# Contact guidance as a consequence of coupled morphological evolution and motility of adherent cells

**DOI:** 10.1101/2021.06.09.447726

**Authors:** Alberto Ippolito, Antonio DeSimone, Vikram S. Deshpande

**Affiliations:** Department of Engineering, Cambridge University, Cambridge CB2 1PZ, UK; SISSA – International School for Advanced Studies, Trieste, Trieste, Italy; The BioRobotics Institute, Scuola Superiore Sant’Anna, Pontedera, Pisa, Italy

## Abstract

Adherent cells seeded on substrates spread and evolve their morphology while simultaneously displaying motility. Phenomena such as contact guidance viz. the alignment of cells on patterned substrates, are strongly linked to the coupling of morphological evolution with motility. Here we employ a recently developed statistical thermodynamics framework for modelling the non-thermal fluctuating response of the cells to probe this coupling. This thermodynamic framework is first extended to predict temporal responses via a Langevin style model. The Langevin model is then shown to not only predict the different experimentally observed temporal scales for morphological observables such as cell area and elongation but also the interplay of morphology with motility that ultimately leads to contact guidance.

**Author Summary:** The evolution of cellular morphology and organization plays a crucial role in the micro-architecture of tissues and dictates their biological and mechanical functioning. Despite the importance of cellular organization in all facets of tissue biology, the fundamental question of how a cell organizes itself in an anisotropic environment is still poorly understood. We demonstrate, using a Langevin style model, that non-thermal fluctuations fuelled by nutrient exchanges between the cell and its environment are critical in allowing cells to explore their surroundings. The biochemical changes, associated with non-thermal fluctuations, drive cell motility and morphological changes and the interplay between these two emerging dynamics ultimately leads to contact guidance, a critical component for tissue morphogenesis.

## Introduction

Living systems are characterised by a wide variety of timescales governing key biological processes at different length scales. For example, on a macroscopic scale it is well known that animal species have their own circadian rhythm [1–2] that governs their daily behaviours, such as feeding and resting. On a microscopic scale cells follow a cell cycle, which is a sequence of cellular phases such growth and division [3–4]. These timescales are critical for cell development and are controlled by complex molecular pathways [3–6]. Another example pertains to the in-vitro spreading of adherent single cells on substrates. Experiments have demonstrated that the spreading and elongation rates differ in time by an order of magnitude [7–8]. The rate of spreading and elongation were also observed to be non constant, which suggested that distinct temporal phases may exist in a cell’s morphological evolution.

Another important timecale pertaining to in-vitro experiments of adherent cells is associated with cell motility. Motility occurs via large and co-ordinated morphological changes in the cell, e.g. the treadmilling mechanism in adherent cells, whereby cells move by tugging on the substrate by exploiting focal adhesions [9–10]. Therefore the motility and morphological change timescales are expected to be interlinked. It is well-known that substrate properties such as stiffness [11–13], chemical composition [13–15] and topology [16–18] strongly affect single cell morphological and motile behaviour. Experiments have shown that changing the substrate stiffness alters both the spreading timescales of adherent cells and the speed at which they explore the environment [19–21]. Guidance provided by substrate anisotropy, i.e. contact guidance [22–24], also influences cell spreading and motile behaviour. In fact, cells elongate more and faster on adhesive channels [25] and that is also accompanied by an enhanced alignment of the biochemical force generating machinery within the cell, viz. stress-fibres [26]. Measurements have also reported a faster directional exploration speed in confined settings [27]. Physical insights into the interplay between morphological changes and motility for cells on anisotropic substrates are lacking in spite of their importance in the understanding of cell guidance.

The aim of this theoretical work is to understand the timescales governing phenomena such as guidance that emerge due to the coupling of morphological evolution with cell motility. We propose a novel framework that we call the Homeostatic Langevin Equation (HLE) that is an extension to the Homeostatic ensemble [28] developed to understand the long time scale or stationary responses of cells. The HLE recognises the non-thermal fluctuating response of the cells and is used to simulate the temporal evolution of isolated cells seeded on unpatterned and patterned substrates. A schematic of the simulation setup for a cell seeded on a substrate with a fibronectin stripe of width *W* is shown in Fig. 1a along with a representative prediction in Fig. 1b. The cell seeded from suspension is circular and the HLE will predict its coupled morphological evolution along with its motility as shown in Fig. 1b. Key experimental observables such as cell area, cell shape, distribution of cytoskeletal proteins such as stress-fibre distributions, shape of nucleus and focal adhesion distributions are outcomes of the simulations. These predictions are used to extract the timescales of morphological evolution and cell motility and thereby develop an understanding of the mechanisms and processes that result in the guidance of cells on patterned substrates.

**Figure 1:**
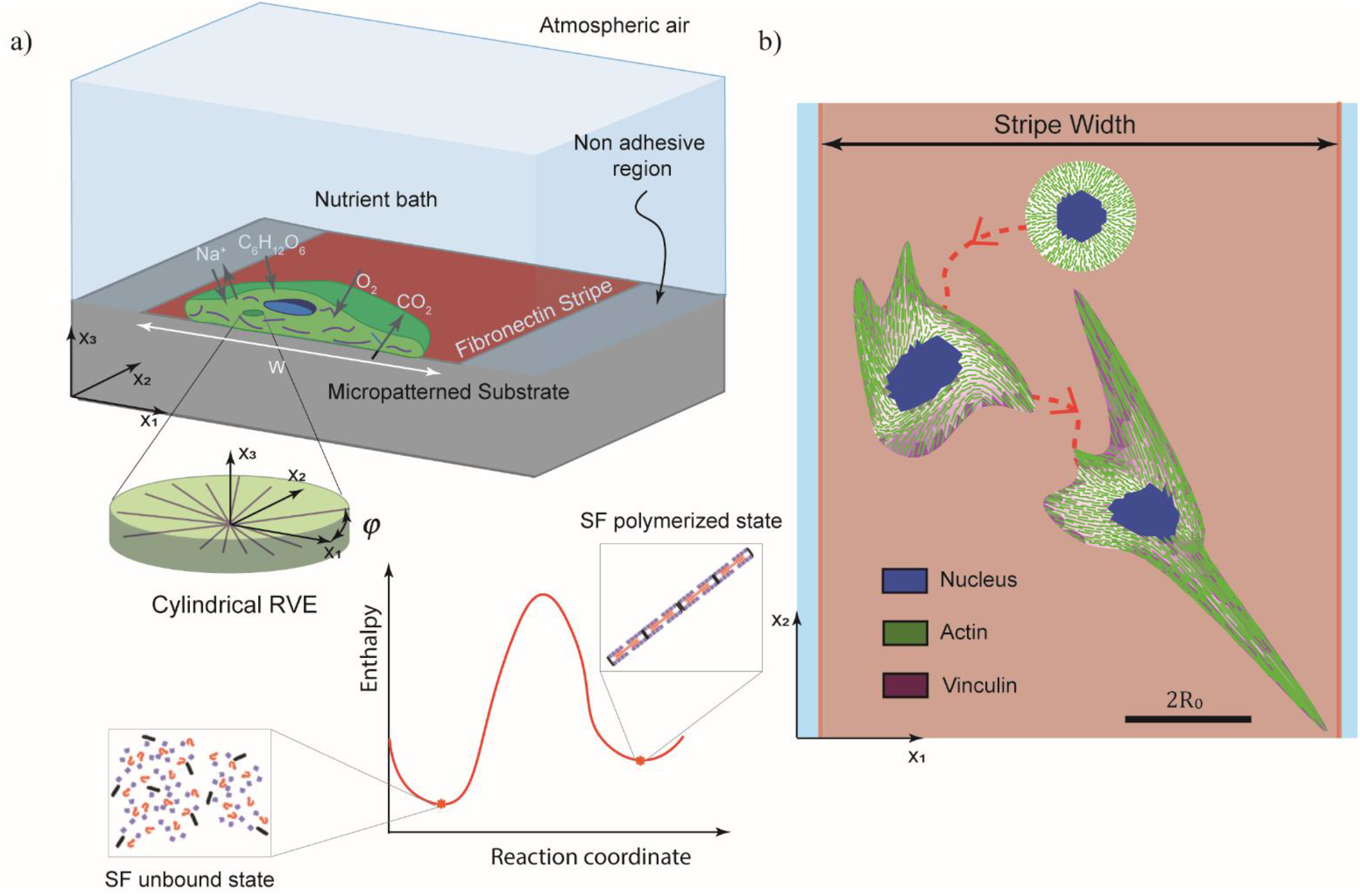
Simulation set-up. (a) Single cell placed on a fibronectin stripe of fixed width *W*. The cell exchanges high energy nutrients with the nutrient bath. The components of the cell modelled explicitly include an elastic nucleus and cytoplasm as well as the contractile stress-fibres in their polymerised state in equilibrium with their unbound components within the cytoplasm. (b) Temporal evolution of an adherent cell on a fibronectin stripe. In the simulation, cells are modelled as two-dimensional bodies in the *x*_1_ – *x*_2_ plane lying on a substrate. All cells are seeded from their state in suspension and then spread and change shape while simultaneously exploring the substrate. The images in (b) are representative images from simulations with the scalebar is 2*R*_0_ the diameter of the circular cell in its elastic resting state.

## Methods

The interaction of cells with enviromental cues (e.g. mechanical, chemical) has a profound influence on cell response. For example, not only do morphological observables such as cell area and aspect ratio increase with increasing substrate stiffness [7–8] but cell motility also exhibits a biphasic behaviour on stiffer substrates [29–30]. While different modelling frameworks are used to understand the influence of environmental cues on cell morphology [31–33] and cell motility [34–35], the interplay of the underlying mechanisms remains elusive. Our aim here is to develop a unified framework with the objective of better elucidating the underlying mechanisms. We do this by extending the recently developed *homeostatic mechanics framework* [28], which predicts the distribution of morphological states that an adherent cell assumes in the interphase period of the cell cycle. This framework, however, lacks temporal information and here we extend the formulation to enable predictions of the temporal evolution of morphological observables alongside cell motility as a function of environmental cues.

### A brief overview of the homeostatic mechanics framework

The homeostatic mechanics framework (further details in [28]) recognises that a cell is an open system which exchanges nutrients with the surrounding nutrient bath (Fig. 1a). These high-energy nutrient exchanges fuel large fluctuations (much larger than thermal fluctuations) in the cell response associated with various intracellular biochemical processes. The cell uses these biochemical processes to maintain itself in the homeostatic state.

Specifically, homeostasis is the ability of a living cell to remain out of thermodynamic equilibrium by maintaining its various molecular species at a specific average number that is independent of the environment [36]. This average number is sustained over all the non-thermal fluctuations of the cell (at-least over the inter-phase period of the cell cycle and in the absence of any imposed shock such as starving the cell of nutrients). The implication is that over the fluctuations of the cell from any reference state, 〈Δ*N_i_*〉 = 0 where Δ*N_i_* is the change in the number of molecules of species *i* from its reference value with 〈x〉 denoting the average of x over the ensemble of states sampled over the non-thermal fluctuations. These fluctuations alter the cell morphology, and each morphological microstate (cell shape, protein distribution etc.) has an equilibrium Gibbs free energy 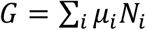, where *μ_i_* is the chemical potential of species *i*. Using the Gibbs-Duhem relation, we then rewrite this in terms of the reference state as 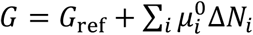, where now 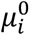 is the chemical potential of species i in the reference state and *G*_ref_ is the equilibrium Gibbs free-energy the cell in its reference state. Upon employing the homeostatic constraint that 〈Δ*N_i_*〉 = 0, we have 〈*G*〉 = *G*_ref_, i.e., irrespective of the environment, the ensemble average Gibbs free energy is constant. This is a universal constraint that quantifies the fact that living cells maintain themselves out of thermodynamic equilibrium but yet attain a stationary state which is the homeostatic state. While the above constraint specifies the average state of the cell, it remains to determine the distribution of states the cell assumes that satisfies this average. As mentioned, cells do not remain in a single microstate but fluctuate between different configuration after seeding. Cells explore these different morphological microstates thanks to several biochemical processes, such as actin polymerisation and treadmilling. The objective of the homeostatic mechanics framework is to capture the different configurations by adopting the *ansatz* that the observed distribution of cell shapes is the one with the overwhelming number of microstates, i.e., the distribution that maximises the *morphological entropy* subject to the homeostatic constraint, i.e. 〈*G*〉 = *G*_ref_ = *G*_S_, with *G*_S_ being the free energy of a cell in suspension, and to any other geometrical constraints such as confinement imposed by adhesive patterning on substrates. The distribution of states is denoted as an equilibrium distribution, where “equilibrium” means in a homeostatic state.

In this study the cells are modelled as two-dimensional bodies in the *x*_1_ – *x*_2_ plane lying on a substrate (Fig S. 1). In this two-dimensional context, the cell morphology is defined by the positional vectors 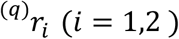 of the *q* = 1,…, *M* material points used to specify the cell shape (Supplementary Section 2.1 for details). Denoting the positions of the material points in morphological microstate (*j*) as 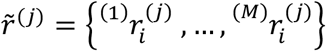 we have the Gibbs free-energy of the cell in microstate (*j*) as 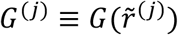. Shishvan et al. [28] demonstrated that the cell in homeostasis assumes a given morphological microstate (*j*) with probability

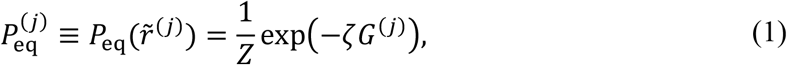

where *Z* ≡ ∑_*j*_ exp(–*ζ G*^(*j*)^) is the partition function of the morphological microstates, and the distribution parameter *ζ* emerges from the homeostatic constraint 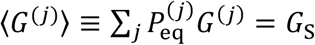. Thus, 1/*ζ* in (1) is referred to as the *homeostatic temperature*, and it sets the equilibrium distribution of morphological microstates of the cell (also referred to as the *homeostatic ensemble*) as an analogous quantity to the thermodynamic temperature of the canonical ensemble [37].

### Gibbs free-energy of a morphological microstate

The implementation of the homeostatic mechanics approach described above requires a specific model for the Gibbs free-energy of the cell-substrate system in a given morphological state. Modelling all the elements of the cell is unrealistic, given that many of their kinetics are still unknown, and often not required, as specific components are known to determine and control the cell response to different environmental cues. Here, we are interested in investigating the response of cells to adhesive patterning of the substrates. These cues are known to guide single cell behaviour resulting in significant cell alignment bias as well as remodelling of the stress-fibre cytoskeleton. Thus, we use a model that includes stress-fibre cytoskeleton introduced by Vigliotti et al [38] and subsequently modified in [26] as well as passive elasticity of the cytoplasm and nucleus. Details of the model including the parameters are given in Supplementary Section 1 and here we give a brief review.

Single cells are modelled as two-dimensional bodies in the *x*_1_ – *x*_2_ plane lying on a substrate such that the out-of-plane Cauchy stress *Σ*_33_ = 0 (Fig S.1). The cell model consists of three essential components: a cytoplasm that is modelled as comprising an active stress-fibre cytoskeleton wherein the actin and myosin proteins exist either in unbound or in polymerised states (Fig. 1a), a passive elastic nucleus, and elements such as the cell membrane, intermediate filaments and microtubules that are all lumped into a single passive elastic continuum. The cell in its undeformed state (also known as the elastic resting state, with the cell elastic strain energy *F*_passive_ = 0 in this undeformed state) is a circle of radius *R*_0_ with a circular nucleus of radius *R*_N_ whose centroid coincides with that of the cell. For a given morphological microstate, the strain distribution within the cell is specified, which directly gives the elastic strain energy of the cell *F*_passive_ via a 2D Ogden-type hyperelastic model for both the nucleus and cytoplasm. The stress-fibre cytoskeleton within the cytoplasm is modelled as a distribution of active contractile stress-fibres such that at each location *x_i_* within the cell, 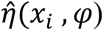 parameterises the angular concentration of stress-fibres over all angles *φ*, while 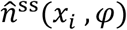 denotes the number of functional units within each stress-fibre. Thus, at any *x_i_* there is a total concentration 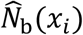 of bound stress-fibre proteins obtained by integrating 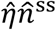 over all orientations *φ*, and these bound proteins are in chemical equilibrium with the unbound stress-fibre proteins (Fig. 1a). The unbound proteins are free to diffuse within the cell, and thus at equilibrium of a morphological microstate, the concentration 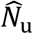 of these unbound stress-fibre proteins is spatially uniform. This chemical equilibrium condition along with the conservation of stress-fibre proteins within the cells provides the spatial and angular distributions of stress-fibres from which the free-energy of the cytoskeleton *F*_cyto_ is evaluated. Then, the total free-energy of the cell-substrate system in morphological microstate (*j*) follows as 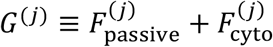.

### Spreading of cells on substrates

It is instructive to briefly review the predictions of the homeostatic mechanics framework prior to extending the framework to predicting temporal evolutions. Recall that the homeostatic framework predicts the stationary distribution of microstates that the cell attains in a given environment. A cell in suspension needs to self-equilibrate and therefore assumes a unique configuration. For the 2D cell modelled here with the choice of fibroblast parameters (Supplementary Table 1), the cell in suspension is a circle of radius 0.92*R*_0_ with the nucleus remaining undeformed with a radius *R*_N_. In this configuration the elastic stresses generated by the compression of the cytoplasm are balanced by the tensile stresses generated by a spatially uniform distribution of stress fibres within the cytoplasm.

Now consider a cell seeded on a patterned substrate as shown in Fig 1b. The substrate is micropatterned with fibronectin stripes of a given width *W* and adhesion of cell is prevented outside the stripes. This theoretical set up is equivalent to experimentally microprinting widely spaced adhesive stripes on the substrates such that the cell cannot span two adhesive stripes and is thus confined to a single stripe. The cell stresses no longer need to be self-equilibrated as stresses within the cell can be balanced via tractions between the cell and the substrate. The cell therefore no longer attains a unique configuration but rather can fluctuate between different morphological microstates with (1) specifying the probability of a specific morphological microstate and the distribution parameter *ζ* set by the requirement that homeostatic constraint is satisfied over the entire ensemble of states the cell can attain. Predictions of the probability distribution of the normalised free-energy *Ĝ* ≡ *G*/|*G_S_*| of the cell are included in Fig. 2a along with the corresponding distributions of the cell area and aspect ratio in Figs. 2b and 2c, respectively. These metrics that are typically used to characterise cell morphology are defined as the normalised cell area *A/A_R_* where *A* and *A_R_* are the areas of the cell and the area of the cell in suspension (*A_R_* = *π*(0.92*R*_0_)^2^), respectively while the cell aspect ratio *A_s_* is defined as the ratio of the major to minor axis of the best fit ellipse. Clearly the cells are predicted to spread and elongate on the substrate with the driving force for these morphological changes arising from the fact that these spread and elongated states have low free-energy: as discussed by Shishvan et al. [28], spreading of the cell reduces its cytoskeletal free-energy but results in an increase in the elastic energy and it is this competition that controls cell spreading. On the other hand, the distribution of morphological states as evidenced in Figs. 2b and 2c is set by the distribution parameter *ζ* and hence by homeostatic constraint. Additionally, we observe that the confinement imposed by the adhesive stripes has a strong tendency to elongate the cells but has a minimal effect on the distribution of the cell free-energy and cell area. These results are consistent with the predictions in [26] and serve as a useful reference as we proceed to understand the temporal evolution of cells seeded on substrates.

**Figure 2:**
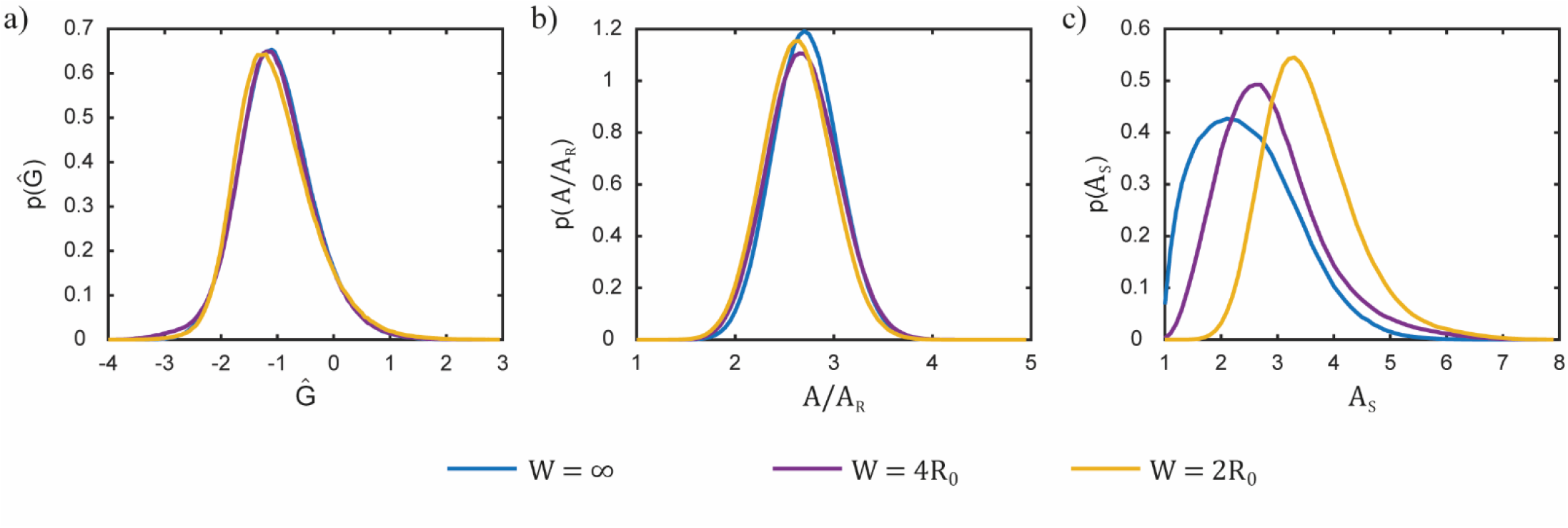
The long-term stationary response of a cell on a rigid substrate patterned with fibronectin stripes of different widths. Predictions of the equilibrium probability distributions of (a) normalised free-energy *Ĝ*; (b) normalised a cell area *A/A_R_* and (c) aspect ratio *A_s_* for stripes of widths of W = ∞, i.e. fully adhesive substrate, 4*R*_0_ and 2*R*_0_.

### A Langevin style framework for cell dynamics

The homeostatic mechanics framework gives the stationary distribution (1) of morphological microstates that the cells will attain in a given environment as seen above. This stationary state is typically attained within 24 to 36 hours after seeding cells into that environment. The framework gives no temporal information about the evolution of the cells as they attain this stationary state including whether all morphological observables reach their stationary distribution at the same rate or whether there are multiple operative timescales. Here we propose the simplest possible extension of the homeostatic mechanics framework to a dynamical setting by invoking an analogy with the canonical ensemble.

Within the context of the homeostatic ensemble, the Gibbs free-energy *G*^(*j*)^ of a cell fluctuates while the cell is in its stationary state (also referred to as homeostatic equilibrium), but the corresponding homeostatic potential 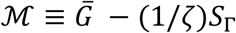 remains constant, where 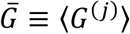 and *S*_Γ_ the morphological entropy of the cell. Thus, a direct analogy can be made between the homeostatic ensemble and the well-established canonical ensemble where the Helmholtz free-energy 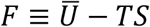 is a constant in a bath with a thermodynamic temperature *T*. In this bath, the system fluctuates over its microstates (*j*) such that it has a fluctuating internal energy *U*^(*j*)^ which at equilibrium achieves average value 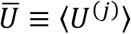 and entropy *S*. Thus, we see that the internal energy *U*^(*j*)^ and temperature *T* in the canonical ensemble are analogous to *G*^(*J*)^ and 1/*ζ*, respectively in the homeostatic ensemble. The temporal evolution of the microstates of an isothermal system whose equilibrium is given by the canonical ensemble is often described by Langevin dynamics. The analogous overdamped (neglecting inertia, due to the low speeds at which cells move [39]) version for the homeostatic ensemble, which we call Homeostatic Langevin Equation (HLE), can be developed as follows. We specify that the temporal evolution of the co-ordinates 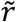 that describe the cell morphology is given by

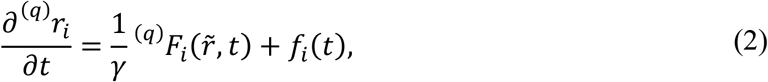

where *t* is time and *γ* is a damping coefficient sometimes referred to as the mobility that relates the velocity of the microvariable 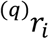 to a determinstic force

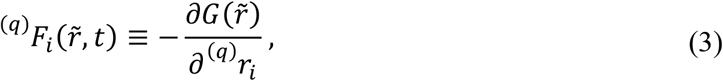

and *f_i_*(*t*) a contribution from a random force. Thus, while the first term on the right-hand-side of (2) represents a determinstic force contribution the second term is the random contribution associated with large non-thermal fluctuations due to high energy nutrient exchanges between the cell and the nutrient bath in which it resides. In principle, *f_i_*(*t*) are correlated in time since the molecular processes from which they originate have a finite correlation times [40]. However, observations [34, 40] suggest diffusive type behaviour of cells over timescales of a few minutes and thus we assume that there exists some correlation timescale *t_c_* over which 〈*f_i_*(*t*)*f_j_*(*t′*)) = *g_ij_*(*t* – *t′*) decays rapidly. Then on timescales ≫ *t_c_* (which in the case of fibroblasts is on the order of a few minutes), *f_i_*(*t*) can be taken to be a delta correlated stationary Gaussian process satisfying 〈*f_i_*(*t*)〉 = 0 and 〈*f_i_*(*t*) *f_j_*(*t′*)) = Γ*δ_ij_δ*(*t* – *t′*) where *δ_ij_* and *δ*(·) are Kronecker and Dirac deltas, respectively and Γ the standard deviation of the random and decorrelated *f_i_*(*t*). Under these assumptions (2) reduces to the Langevin equation.

It now remains to set the standard deviation Γ. In order to set Γ we turn to the Fokker-Planck equation corresponding to the HLE. In the HLE, we expressed the uncertainty in the morphology of the cell in terms of Gaussian correlation functions. Shifting perspective, we can ask: what probability distribution 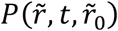 where 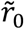 is 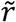 at time *t* = 0 would give the same correlation functions? It is important here to stress that we do not care about that path 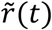 the cell took but rather ask the simpler question of the probability that the cell attains a morphological microstate 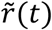 at time *t*, regardless of how it got there. This is equivalent to saying that if we run a large number of independent but nominally identical experiments then what is the probability distribution of morphological microstates observed at time *t*. This is a well-established problem and following Ichimaru [41], the required probability distribution can be shown to satisfy the Fokker-Planck equation

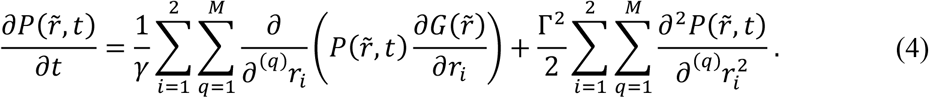

The steady-state solution to (4) corresponding to 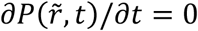 is the equilibrium probaility distribution and given by:

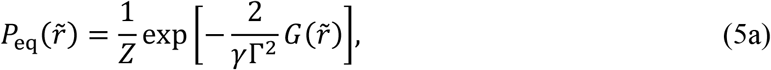

where:

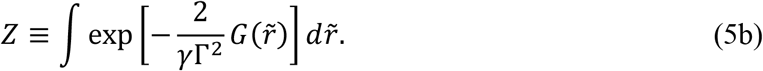

Therefore the Fokker-Panck equation (4) coverges to the homoeostatic ensemble (1) by setting:

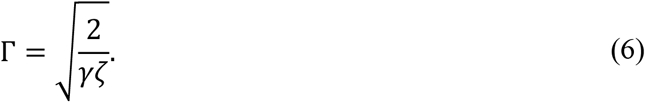

We shall thus use this choice of Γ to resolve the temporal response of a single cell in a given environment. Recalling that *f_i_*(*t*) in (2) is random and decorrelated with a standard deviation Γ we can rewrite (2) in normalised form as:

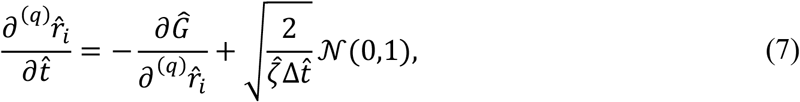

where 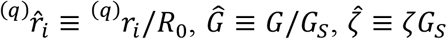 and 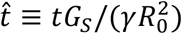 while 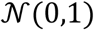 is a Gaussian distribution of zero mean and unit variance. In writing (7) we have used (6) and the fact that the stochastic differential equation (7) is solved with a finite time step Δ*t* with 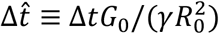. It is now apparent that while *ζ* sets the fluctuation magnitude, the mobility *γ* sets the evolution timescale. We emphasize that this single timescale for the evolution of the morphological microstate does not imply that all observables evolve at the same rate as we shall proceed to show. Further details of the numerical implementation scheme are provided in Supplementary Section 2.3 with the Supplementary videos giving a better sense of the dynamical output of the simulations. All results are presented for model parameters representative of myofibroblasts as calibrated in [26] but we emphasize that the model holds more generally for any adherent single cell type.

## Results

We shall discuss two cases: (i) cells on unpatterned substrates where there is effectively no confining effect imposed by the substrate within the *x*_1_ – *x*_2_ plane and then proceed to contrast with the case of (ii) contact guidance on substrates patterned with adhesive stripes of width *W* (Fig. 1b).

### Cell area and elongation evolve at different rates on unpatterned substrates

We conceive of a typical experiment where a cell in suspension is seeded on a rigid substrate coated uniformly with an adhesive protein such as fibronectin. The evolution of the cell morphology is then observed as a function of time *t* with *t* = 0 corresponding to the instant of seeding. Of course, motility of the cell and the evolution of its morphology are coupled but first we focus on cell morphology. Representative images of computed cell morphologies and the corresponding actin, nucleus and focal adhesion organisations are included in Fig. 3a at three normalised times 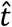 in addition to the state at 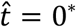^*^. Recall that the Langevin style dynamical equation (7) is a stochastic differential equation so that a different solution is generated for every realisation of the noise process, i.e., much like in experiments a different trajectory of morphological evolution is obtained for every solution of (7) with the same initial state at *t* = 0. Hence in Fig. 1a we show solutions at three times from three such trajectories. While of course the three morphologies computed using different trajectories of (7) are different they show many similar features. These include the observations that with increasing time: (i) the cells spread and increase their area as well as their ellipticity or aspect ratio and (ii) the level of actin polymerisation and focal adhesions also increase.

**Figure 3:**
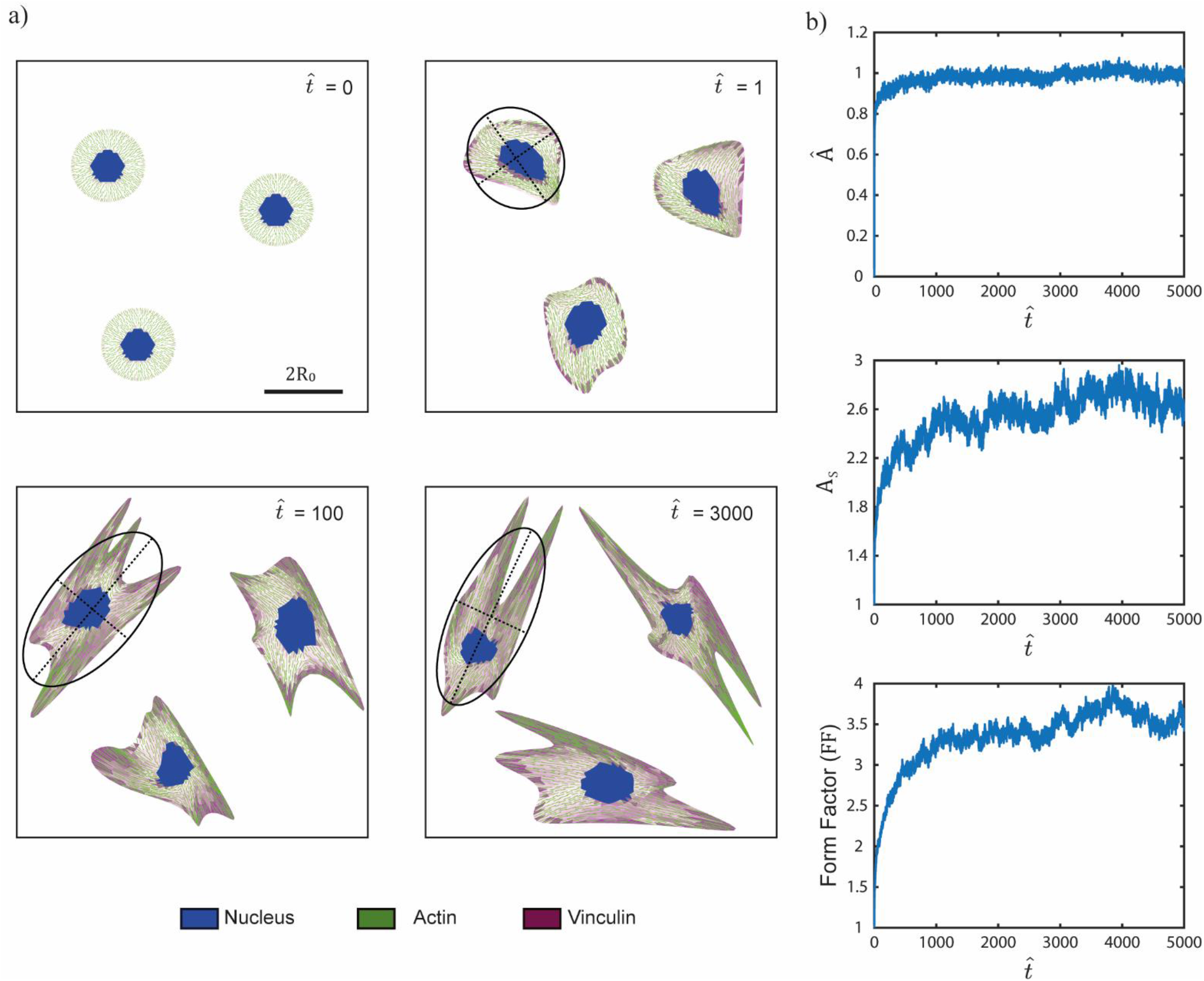
Temporal evolution of cells on unpatterned substrates. (a) Three representative images of cells at three normalised times 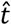 for cells seeded at time 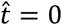. An example of a best fit ellipse is drawn on one of the selected temporal evolution. The scalebar is 2*R*_0_. b) Temporal evolution of morphological observables, viz. normalised cell area *Â*, aspect ratio *A_s_* and form factor FF. The results are shown as averages over *n* = 100 Langevin trajectories.

It is instructive to make quantitative predictions of some morphological observables to enable more direct comparison with measurements. The complete morphology of the cell in the model is decribed by 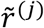 but as in observations [26] we focus on three coarse-grained metrics of the morphology viz. (i) cell area *A*; (ii) cell aspect ratio *A_s_* (defined from a best fit ellipse) and (iii) the form factor FF ≡ *p*^2^/(4*πA*) where *p* is the cell perimiter (the definition of the form factor is chosen such that FF = 1 for a circular cell and rises as the cells forms protrusions such as filopodia). Each solution of the HLE (7) produces a different trajectory and much like in experiments trends are most clearly seen by examining the ensemble average over a number of trajectories. We performed *n* = 100 independent simulations and defined the ensemble average as follows. At time *t*, the ensemble average area is defined as 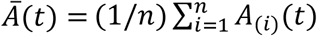, where *A*_(*i*)_(*t*) is the area of the cell in the *i*^th^ trajectory at time *t*. The ensemble average aspect ratio *Ā_s_*(*t*) and 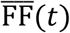 are defined in an analogous manner. While *A_s_* and FF are non-dimensional it is instructive to define a non-dimensional area as

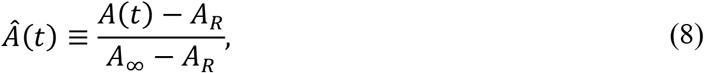

where *A*_∞_ ≡ Ā(*t* → ∞) so that *Â* assumes values of 0 when the cell is seeded and fluctuates around 1 at convergence. Predictions of the temporal evolution with normalised time 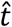 are included in Fig. 3 and are consistent with the qualitative features seen in images of the cell morphologies (Fig. 3a). However, an intriguing feature emerges: while the HLE has only a single time-scale the normalised cell area *Â* evolves and reaches its steady-state much faster compared to the cell aspect ratio and form factor, i.e. different timescales for the evolution of area and shape emerge from the HLE. Also see Supplementary Video 1 for the simulated evolution of the cell over a single trajectory shown alongside the plots of the evolution of the ensembled averaged observables.

The different time-scales for the evolution of cell area and aspect ratio have also been observed in experiments. In Figs. 4a and 4b we reproduce measurements from [7–8] of the temporal evolution of normalised area and aspect ratio for the spreading of fibroblasts on stiff substrates with *t* = 0 corresponding to the instant of seeding the cell from suspension. In Fig. 4a the average cell area over about 50 measurements is plotted with the error bars corresponding to the standard deviation, while in Fig. 4b the average aspect ratio is plotted over 10 measurements (two measured trajectories are also included to give an indication of the large variability in this metric over different trajectories). The striking difference between Figs. 4a and 4b is that while the steady-state cell area is achieved in about 150 mins the cell aspect ratio evolves significantly more slowly such that is is unclear whether a steady-state aspect ratio is achieved 20 hours after seeding. We superimpose on Fig. 4 our corresponding predictions (averaged over 100 different trajectories) with the time-scale of in the simulations chosen to be 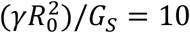 min so as to match the simulated and measured time-scales of the temporal evolution of cell area. With this choice of the time-scale we see that the simulations capture the observation that the cell aspect ratio evolves signficantly more slowly. Moreover, the simulations not only capture the average temporal evolution response but also agree with measurements of the standard deviation of cell area. In addition, two selected simulated aspect ratio trajectories are included in Fig. 4b and illustrate that, in line with measurements, the simulations also display a very large variability of this metric over different trajectories.

**Figure 4:**
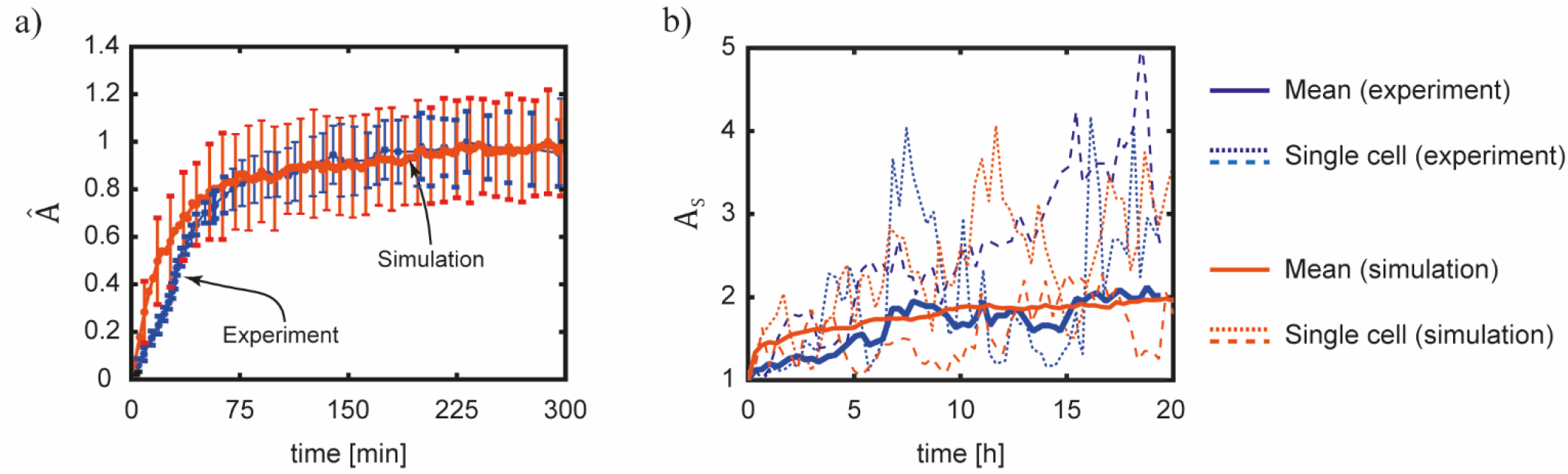
Timescales for the evolution of cell morphology. (a) Comparsion between measurements [7–8] and predictions of the temporal evolution of normalised cell area *Â*. Solid lines and error bars indicate the average and standard deviation, respectively over *n* = 50 measurements and *n* = 100 Langevin trajectories of the simulations. (b) Corresponding comparisons between measurements and simulations of the cell aspect ratio *A_s_*. The solid lines are the average over = 10 measurements and *n* = 100 Langevin trajectories of the simulations while the dotted lines show selected trajectories in the measurements and simulations to indicate the wide variability in both the measurements and simulations. The simulations use a timescale 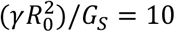 min. This single timescale is shown to capture the obervation that the cell area evolves in minutes while the cell aspect ratio evolves over a timescale of many hours.

The success of the simulations in capturing the fact that the two morphological ovservables evolve at different rates suggests that a single rate constant 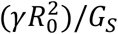 suffices to set the evolution of these observables. We therefore use the model to interogate the source of the two timescales that set the evolution of cell area and aspect ratio. For this we consider a significantly simplified model where the cell is restricted to remain a spatially uniform ellipse with a fixed orientation (Fig. 5a). In this case the stretches *λ*_1_ and *λ*_2_ of the principal axes of the ellipse completely define the cell morphology rather than the *M* positional vectors 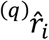. The determinstic evolution of the cell morphology in this simplied model is then given by an equation analogous to (7) with the noise term neglected, viz.

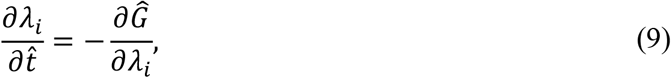

where *i* = 1, 2. Predictions of the temporal evolution of *Â* and *A_s_* (where cell area *A* = *Λ*_1_*Λ*_2_*A_R_* and *A_s_* = *λ*_1_/*λ*_2_ with *λ*_1_ ≥ *λ*_2_) are included in Fig. 5b for a cell seeded from suspension at 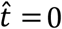. Intriguingly, the qualitative feature that the cell area attains its steady-state significantly faster than aspect ratio is retained in this very simplistic setting. We observe three temporal regimes: (i) an intial regime I of rapid cell spreading where the cell remains circular with *A_s_* = 1; (ii) a subsequent regime II of cell elongation where cell area is constant but the aspect ratio increases and (iii) a final regime III where both cell area and aspect ratio increase although the changes in this regime are relatively minor. Regimes I and II set the two observed timescales for the evolution of cell area and apsect ratio, respectively. In this deterministic and simplistic setting where the cell morphology is only a function of (*λ*_1_, *λ*_2_) we can understand this by examining the energy landscape *Ĝ*(*λ*_1_, *λ*_2_) shown in Fig. 5c. We include in Fig. 5c the trajectory the cell takes in (*λ*_1_, *λ*_2_) space starting from its state in suspension along with isolines of *A/A_R_* = *λ*_1_*λ*_2_ and *A_s_* = *λ*_1_/*λ*_2_. The trajectory set by Eq. (9) has two distinct branches: (i) an initial branch corresponding to regime I where the cell traverses a path of *A_s_* = 1 while *A* increases and then turns and traverses a path of constant *A* but with increasing *A_s_*. This trajectory is purely set by the topology of the *Ĝ*(*λ*_1_, *λ*_2_) landspace and the fact that (9) requires the cell moprhology to evolve along a path with the steepest gradient in *Ĝ*(*λ*_1_, *λ*_2_) space. Thus, we argue that the two different time-scales for the evolution of cell area and aspect ratio are purely a result of the free-energy landscape of the cell. This landscape is set by the interplay between the elastic energy and cytoskeletal energy of the cell; see Supplementary Section 1 for a summary of the free-energy model used here.

**Figure 5:**
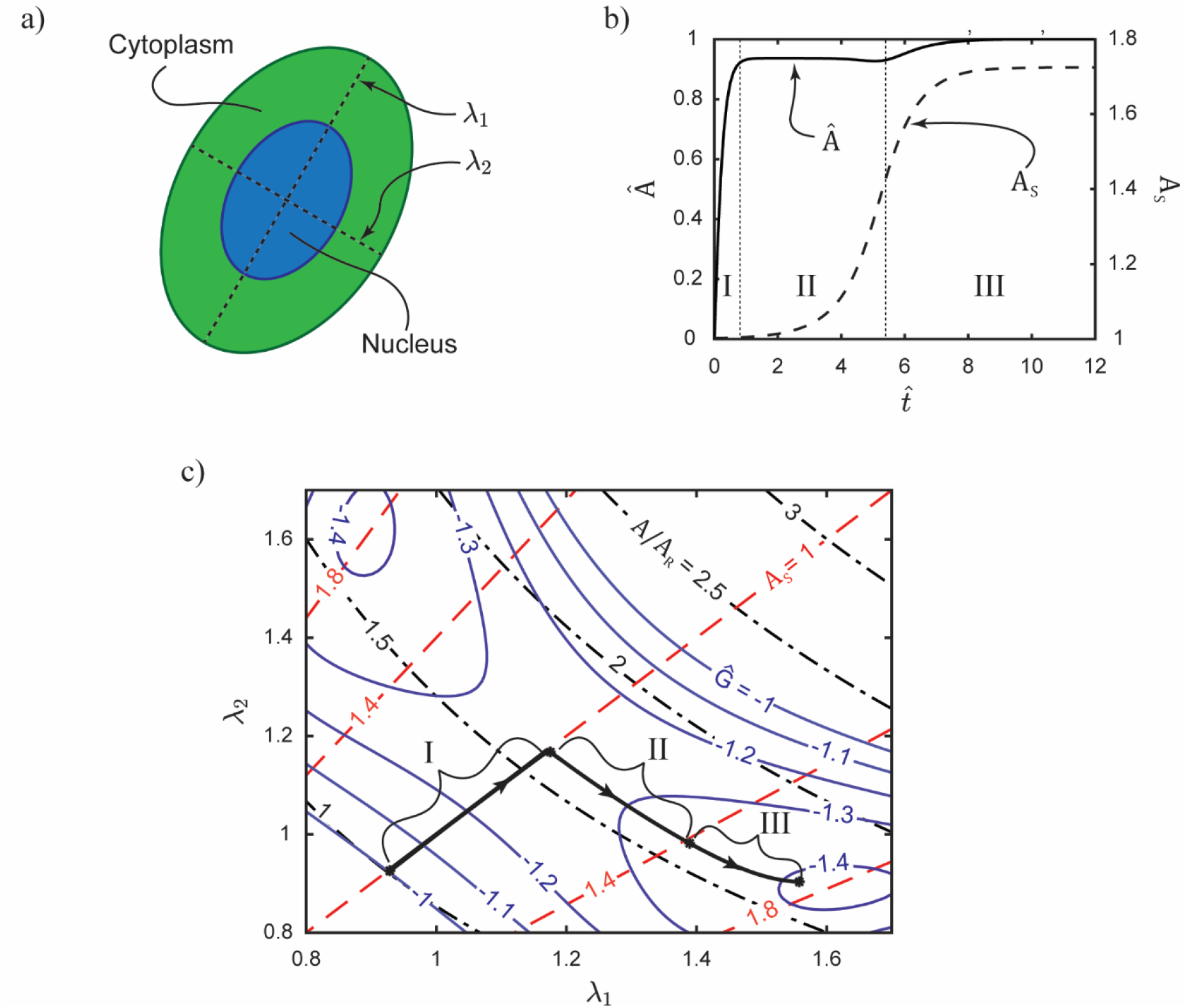
A reduced model of a cell as a spatially uniform ellipse to illustrate morphological evolution. (a) Sketch of the cell including its nucleus as a spatially uniform ellipse. Here *λ*_1_ and *λ*_2_ are the stretches of the axes of the ellipse. (b) Predictions of the temporal evolution of the cell area *A* = *λ*_1_*λ*_2_*A_R_* and aspect ratio *A_s_* ≡ *λ*_1_/*λ*_2_ (*λ*_1_ ≥ *λ*_2_) for a cell seeded on an unpatterned substrate from suspension at time 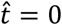. The predictions are for a deterministic response with effects of biological noise excluded. (c) Free-energy landscape of the reduced cell model. Contours of the normalised free-energy *Ĝ* are included on map with axes *λ*_1_ and *λ*_2_ along with contours of *A/A_R_* and *A_s_*. On the map we show the deterministic trajectory of the cell in the free-energy landscape for a cell starting from its state in suspension until it attains its mininum free-energy state on the substrate.

### Diffusive motility of cells on unpatterned substrates

When cells are seeded on substrates coated with an adhesive protein not only does their morphology evolve but this morphological evolution is coupled to their motility. The HLE (7) enables predictions of this coupled moltility and mophological evolution of the cell. Predictions of the coupled evolution of the cell morphology and its motility are are shown in Fig. 6 for 4 selected trajectories of (7) with the cell seeded from suspension at same location at time 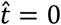 in each case. Stochastic motility (i.e. cells take a random path over the substrate surface) is predicted over the timescales in Fig. 6a in line with numerous observations [34, 42–45]. The sochastic motility can be characterised in terms of the squared displacement

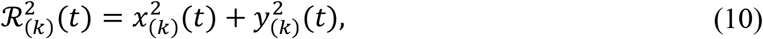

of the centroid of cell (*k*) from its seeding location (taken to be the origin of the Cartesian co-ordinate system with *x*_(*k*)_(*t*) and *y*_(*k*)_(*t*) the Cartersian co-ordinates of the cell centroid at time *t*). Predictions of 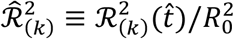 are included in Fig. 6b for 5 trajectories: we predict large variations in 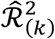 over different trajectories consistent with observations [34]. (The time-lapse of the motility of a single trajectory of a cell along with the corresponding 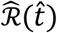 is included in Supplementary Video 2). Given this stochastic nature of the motility it is more instructive to consider the mean squared displacement 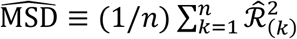 over *n* independent trajectories. The predictions of 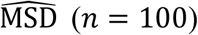 are included in Fig. 6b and illustrate the diffusive nature of the predicted motility as 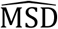 scales linearly with normalised time 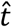. If the cell is assumed to be a single particle whose position is given by the cell centroid, the HLE (7) would predict a diffusion co-efficient *D* = 1/(*γζ*) such that the slope of the 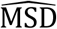 curve in Fig. 6b is given by the normalised diffusion co-efficient 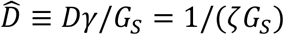. The 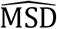 slope in Fig. 6b ≈ 0.06 while for the cell parameters used here 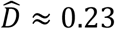, i.e. the actual motility of the cell as predicted by (7) is significantly more sluggish compared to modelling the cell as a Brownian particle with a diffusion co-efficient 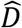. The slower motion is related to the morphological evolution of the cell that is coupled to its motility. If the cell were modelled as a single Brownian particle all fluctuations would result in motion of the particle. On the other hand, in (7), the fluctuations affect the positional vectors 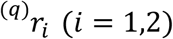 of the *q* = 1,…, *M* that describe the cell morphology and a displacement of the cell centroid requires the co-ordinated motion of these vectors. Such co-ordinated motion via stochastic fluctuations occurs with a low probability resulting in the effective diffusion co-efficient of the cell centroid being significantly smaller than 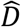.

**Figure 6:**
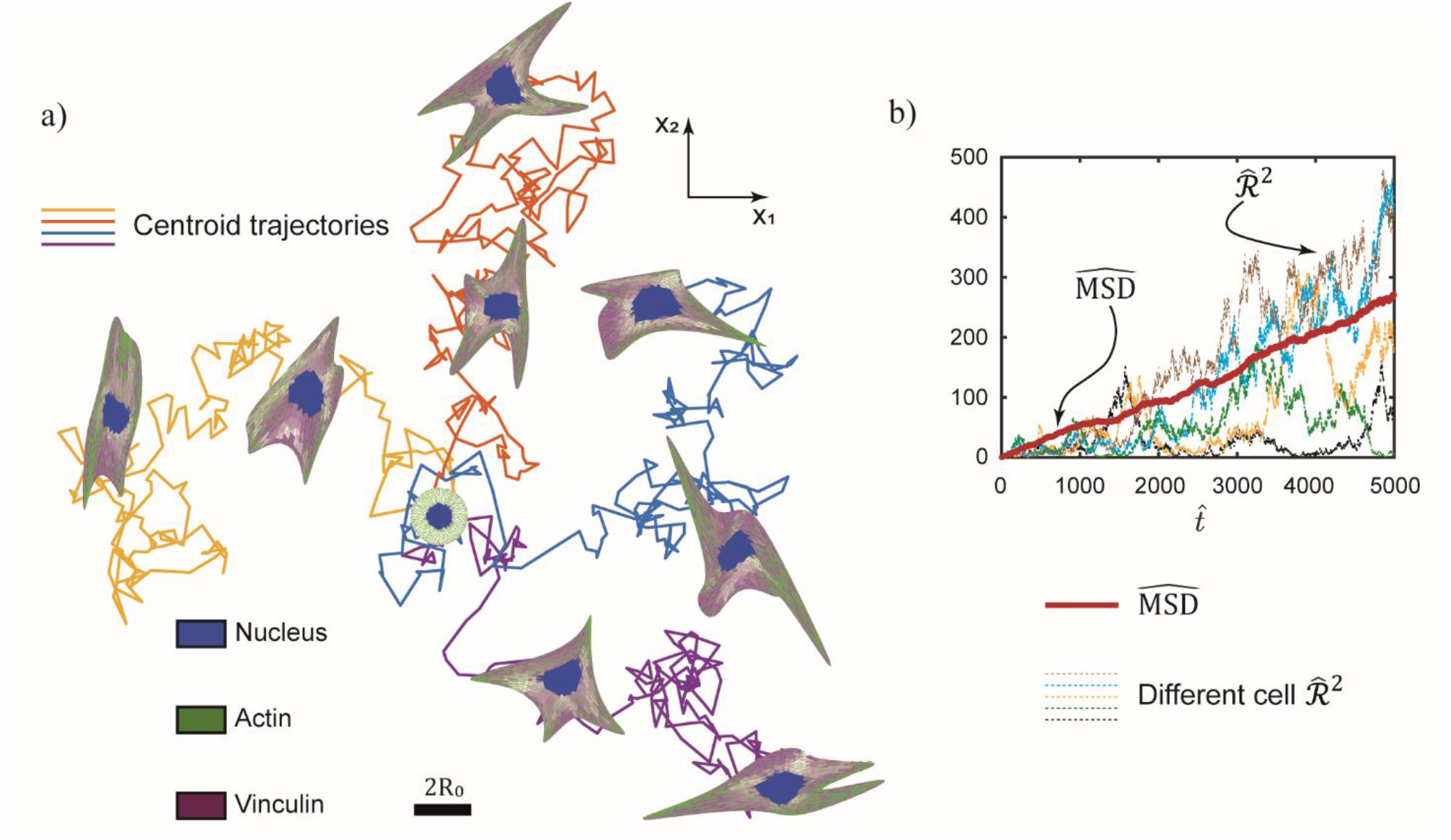
Coupled motility and morphological evolution of the cell seeded on an unpatterned substrate. (a) Four Langevin trajectories for a cell seeded at time 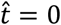 from suspension on an unpatterned substrate. The scalebar 2*R*_0_ is the diameter of the circular cell in its elastic rest state. (b) Predictions of the corresponding temporal evolution of the normalised mean square displacement 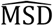 over *n* = 100 Langevin trajectories. Four indivual trajectories parameterised by their normalised squared displacement 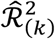 are also included to illustrate the variability in the trajectories as seen in (a).

Finally, we emphasise that our model captures the coupled evolution of cell morphology and motility over time-scales ≫ *t_c_*, i.e. we only capture the diffusive motion of the cell and not its ballistic motion [43] due to persistence in the direction of cell motion over time-scales ≪ *t_c_*. A consequence of the model only capturing the diffusive part of the motion is that it does not provide insights into the physical processes, such as actin treadmilling, that give rise to ballistic motion [40, 46–48].

### Contact guidance on substrates with adhesive stripes

*In vivo*, adherent cells interact with the surrounding extracellular matrix (ECM) that is not only responsible for the structural integrity of tissues but also establishes and maintains the cellular microenvironment by providing cells with mechanical, biochemical, and physical cues. In particular, it is now well-established that cellular microenvironments induce cells to align and migrate along the direction of the anisotropy—a phenomenon called contact guidance [17]. Various *in-vitro* chemical micropatterning approaches using two-dimensional (2D) substrates have been developed to study cellular contact guidance, as model systems to simplify the highly complex three-dimensional (3D) environments *in vivo* [26]. A common micropattern is fibronectin stripes of a given width *W* with adhesion of cell prevented outside the stripes [16, 26]. We now proceed to investigate via the HLE the effect of the confinement provided by the stripes on the evolution of cell morphology and the accompanying motility which finally leads to contact guidance.

Given a cell seeded in the middle of the stripes at time 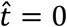 from suspension, Fig. 7a shows snapshots of the cell morphology and position from a single trajectory on stripes of normalised width 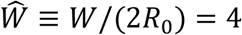 and 1 as well as the unpatterned substrate with 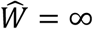 (distribution parameter *ζ* varies with stripe width and the homeostatic ensemble predictions of the variation of *ζ* with 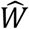 are included in Fig. S2). The snapshots are shown for two selected times along with best fit ellipses and the corresponding orientations *ϕ* of the cell with respect to the stripe. From the *n* ≥ 50 trajectories computed here we have selected in Fig. 7a trajectories where the cell motion backed onto itself between times 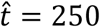 and 1500 so that the cell positions are well separated, and we can easily illustrate the cell morphologies. While clear visual differences between the cell morphologies are observed for the cell on the different stripes, the most striking differences are in the cell trajectories where we clearly see that the cell is “guided” on the 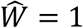 stripe with the trajectory being nearly one-dimensional (also see Supplementary Video 3). This guidance can be characterised in two ways: (i) via the cell orientation that is related to cell morphology and (ii) via the direction of motion. Here we first focus on cell orientation and in the following section on guidance which is more directly a consequence of motility.

**Figure 7:**
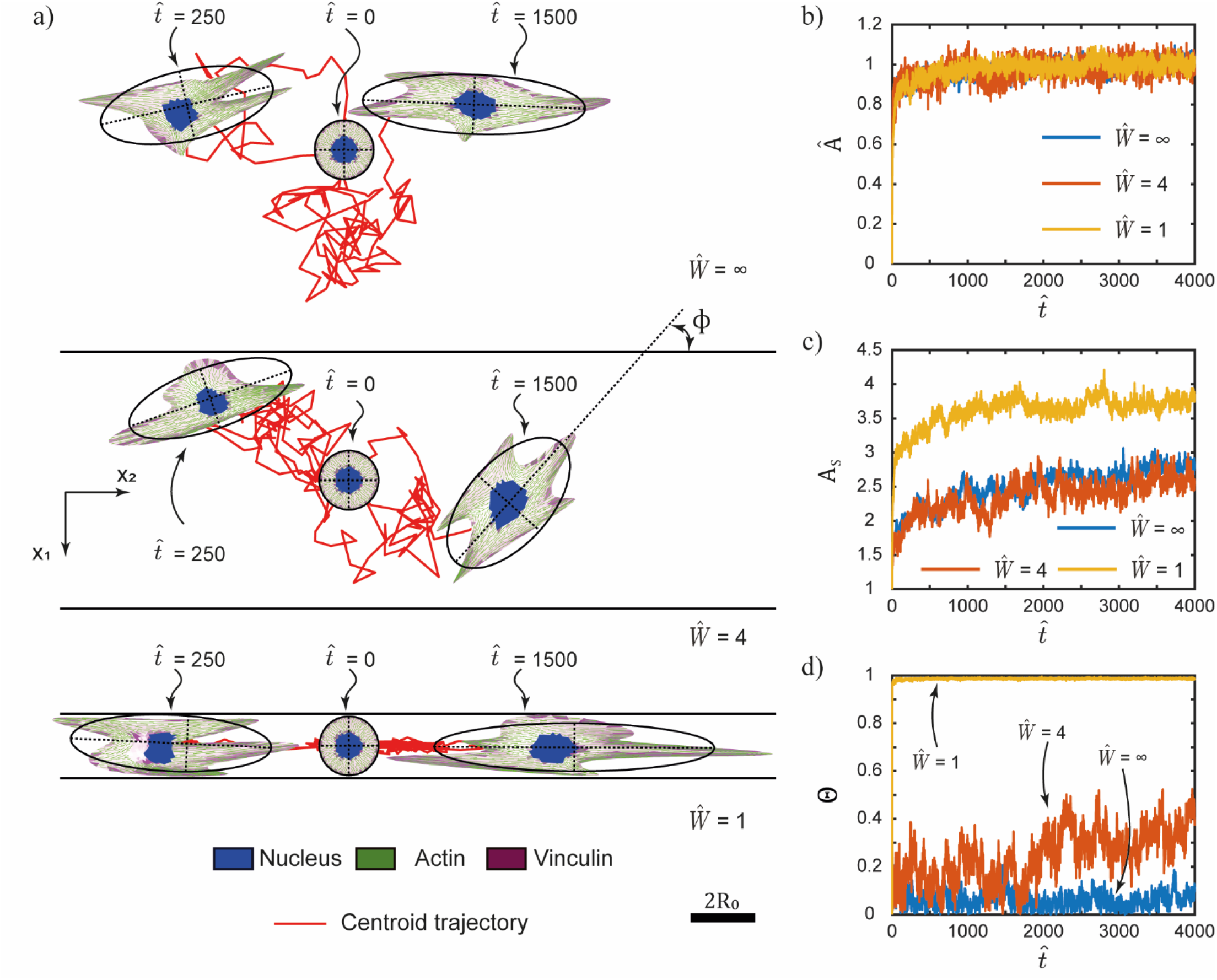
Coupled motility and morphological evolution of the cell seeded on a substrate with adhesive stripes. (a) Two Langevin trajectories for a cell seeded at time 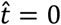 from suspension on a a substrate with adhesive stripes of normalised width 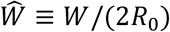. One trajectory shows the evolution upto time 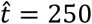 while the second to 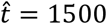. Results are shown for three stripe widths 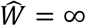, 4 and 1. The scalebar 2*R*_0_ is the diameter of the circular cell in its elastic rest state. The corresponding temporal evolution of (b) the normalised cell area *Â*, (c) aspect ratio *A_s_* and (d) order parameter Θ averaged over *n* ≥ 50 Langevin trajectories.

To characterise the changes in cell morphology with stripe width we plot in Figs. 7b–7d the temporal evolution of the normalised area *Â*, aspect ratio *A_s_* and the order parameter Θ defined as

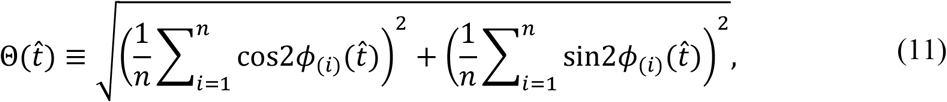

where *ϕ*_(*i*)_ is the orientation of the cell in the *i*^th^ trajectory at time 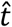 (see Fig. 7a for the definition of *ϕ*_(*i*)_ as the inclination of the major axis of the best fit ellipse to the stripe direction). The order parameter Θ is defined such that Θ = 0 if *ϕ*_(*i*)_ is uniformly distributed over all *n* trajectories while Θ = 1 if *ϕ*_(*i*)_ takes a unique value. Cell area and its temporal evolution are insensitive to 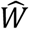 but when the stripe width is on the order of the cell size, i.e., 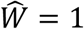, the cell aspect ratio attains a higher steady-state value. Similarly, cell alignment as parameterised through Θ also increases with decreasing 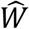 although the large increases in Θ occur at around 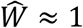 when the aspect ratio too increases. The increase in cell alignment at steady-state with decreasing 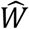 has been reported in [26]. In that study the authors argue that alignment results from the fact that cells near the edge of the stripes are necessarily aligned with the stripes: this boundary effect is of course more prominent for narrower stripes and also with cell elongation. As a consequence, cells are more aligned for smaller 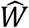 in line with the HLE predictions. Since alignment is a result of cells sensing the stripe edges, it follows that alignment is a consequence of cells wandering over the stripes including towards the edge of the stripes and hence an outcome of cell motility. We therefore expect that the timescale for the evolution of Θ to be set by the motility timescales of cells on the stripes.

### Time-scale for the development of guidance and non-diffusive motility of cells on substrates with adhesive stripes

To quantify the motility of cell on stripes we include predictions of the variation of 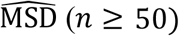 with 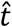 on stripes of different widths in Fig. 8a. Unlike the case of the unpatterned substrate 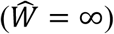, the 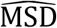 on stripes of width similar to cell size (i.e., the 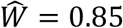 and 1.5 cases shown in Fig. 8a) does not seem to vary linearly with 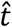. To understand the effect of the constraint of the stripes on motility we introduce a measure of the mean square displacement in the direction of the stripes, viz.

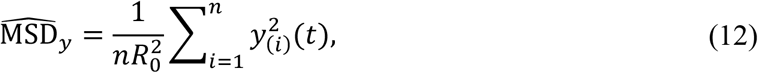

where the seeding location at time *t* = 0 is assumed to be the origin of the co-ordinate system (*y* – aligned with the stripe direction as shown in Fig. 7a). To parameterise the influence of the constraint of the stripes we then define

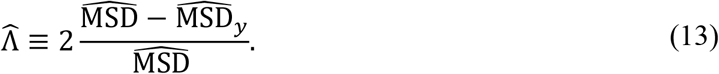

**Figure 8:**
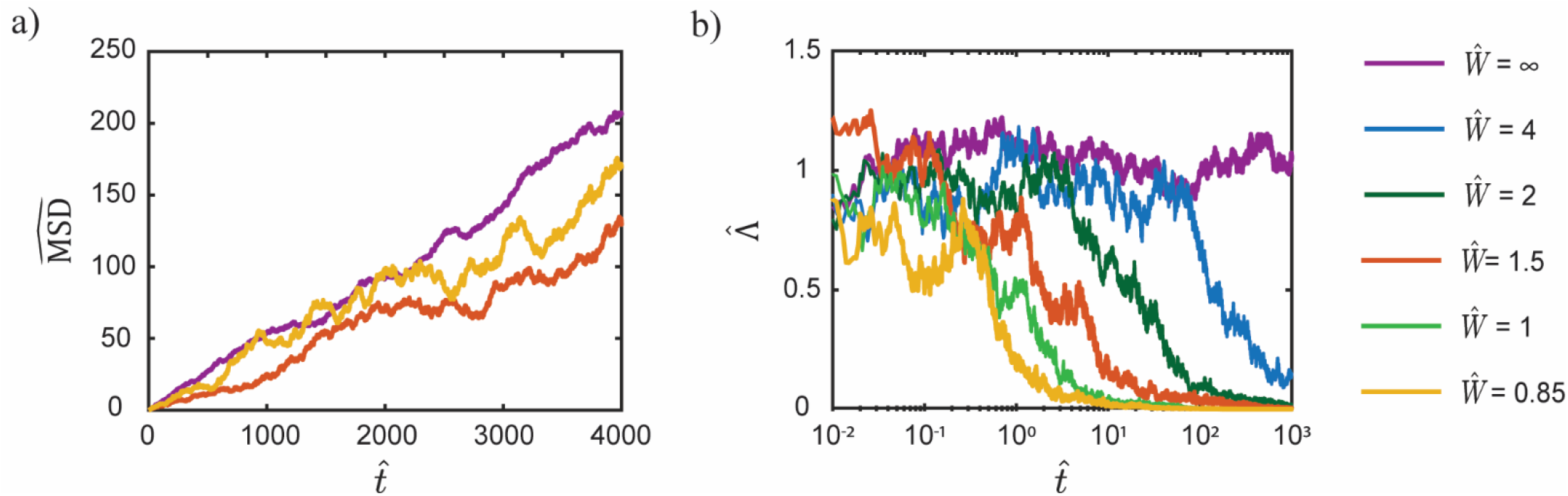
Motility of cells on substrates with adhesive stripes. Predictions of the temporal evolution of (a) normalised mean square displacement 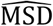 and (b) dimensionality 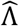 of the motility on stripes of normalised width 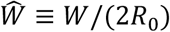. The cell is seeded from suspension at time 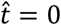 on the stripes.

This parameter that characterises the dimensionality of the motility is defined such that 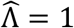 on an unpatterned substrate as 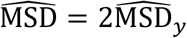 since movements in the *x* and *y* −directions are free and non-distinguishable while 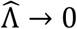 for 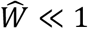 when the motion of the cell is completely constrained to be only in the *y* −direction (i.e. is one-dimensional) so that 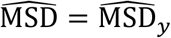. Predictions of the temporal variation of 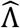 are included in Fig. 8b for selected stripe widths 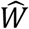. In all cases, 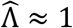 at early stages of the cell motion but, except for the unpatterned substrated 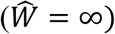, 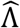 subsequently reduces to 0. However, the time at which this transition from 2D to 1D motility occurs increases with increasing 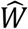. This can be understood by recognising that the *x* −displacement of the cell is constrained by the stripe width while the y −direction displacement is unconstrained and thus with increasing time 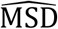 is dominated by the *y* −direction displacement and 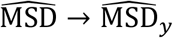. The transition from a two-dimensional (2D) motion of the cell during the early stages of cell motion to 1D motility in the later stages (with the transition time being 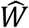 dependent) induces the loss of the linear dependence of 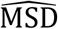 with time seen in Fig. 8a for the finite stripe widths. Moreover, we would also anticipate that the attainment of the steady-state of the order parameter Θ is governed by the time of transition from 2D to 1D motility. A comparison of Fig. 7d with 8b confirms that indeed Θ attains it steady-state value at approximately the time that 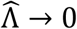 and thus the time to achieve steady-state alignment decreases with decreasing 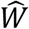. Finally, we note that the transition of 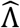 from 1 to 0 denotes the guidance of the cells by the stripes with the time taken for 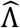 to transition from 1 to 0 the time for contact guidance to be achieved on the patterned substrate.

## Concluding discussion

The temporal response of isolated cells on unpatterned and patterned substrates has been investigated via a novel framework, labelled the Homeostatic Langevin Equation (HLE), that recognises the non-thermal fluctuations of cells. The morphological evolution is driven by gradients of the cell free-energy and we show that the HLE correctly predicts that the cell area or spreading evolves at a rate an order of magnitude faster than cell aspect ratio or elongation. These two very different timescales are not associated with different intracellular molecular timescales but rather just an emergent outcome of the topology of the free-energy landscape of the entire cell. The framework enables the prediction of the coupled evolution of cell morphology along with cell motility. Over the timescales when the HLE is applicable, the simulated cell motility on an unpatterned substrate is Brownian, in line with numerous obervations [34, 42–43], and emerges from coordinated morphological fluctuations.

On substrates patterned with adhesive stripes cells again spread and elongate much like on unpatterned substrates. However, the Brownian motion of the cells is now restricted ouside of the parallel direction to the stripes. The HLE then predicts that if sufficient time is given for the cells to explore the stripe widths their 2D Brownian motion essentially reduces to one-dimensional (1D) motion along the length of the stripes. This results in an apparent non-brownian motion of the cell with the mean-square-displacement of the cell centroid no longer scaling linearly with time over the entire duration. A more important consequence of this switch to 1D motility is contact guidance or rather 1D motion of the cell along the stripes which manifests itself also in terms of the alignment of the cell orientation with the stripe direction. Thus, a key conclusion of the HLE framework is that the non-thermal cell fluctuations give the cell its ability to explore the stripe width and in turn, rather counterintuively, result in its guidance and alignment with the anisotropy of its environment.

## Acknowledgements

A.I. acknowledges support from the Engineering and Physical Sciences Research Council. A.D. acknowledges support from the European Research Council through European Research Council Advanced Grant 340685-Micromotility.

## Author contributions

A.I. performed the simulations, processed the numerical data and created the figures. V.S.D. designed and supervised the project, aided by insightful discussions with A.D. V.S.D and A.I. wrote the manuscript. All authors provided critical comments and edited the manuscript.

## SUPPLEMENTARY

### 1. Free energy of the 2D cell

Here we simplify the 3D cell on the substrate as shown in Fig. S1 as a 2D body of thickness *b*_0_ in the elastic rest state. Here we briefly describe the free-energy model for this 2D cell. With the system comprising of the cell and the rigid substrate within a constant temperature and pressure nutrient bath, the Gibbs free-energy 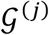 of the system in morphological microstate (*j*) is given by 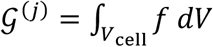, where *f* is the specific Helmholtz free-energy of the cell. Here, there is no contribution from the substrate as it is assumed to be rigid. We emphasize that the analysis presented here is for the system under atmospheric pressure conditions. Therefore, without loss of generality, we set the pressure equal to zero (i.e. use gauge pressure), and thus a pressure term does not appear in the expression for 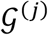. The equilibrium free-energy *G*^(*j*)^ is then the value of 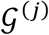 at 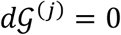. In the following, for the sake of notational brevity, we shall drop the superscript (*j*) that denotes the morphological microstate, as the entire discussion refers to a single morphological microstate.

The cell model is based on the work of Vigliotti et al. [1] and the subsequent modifications of Shishvan et al. [2] and Buskermolen et al. [3]. The model assumes only three elements within the cell: (i) a passive elastic contribution from elements such as the cell membrane, intermediate filaments and microtubules, (ii) an active contribution from contractile acto-myosin stress-fibres that are modelled explicitly and (iii) the nucleus modelled as a passive elastic body. We shall first describe the modelling of the active acto-myosin stress-fibres in the cytoplasm and then discuss the elastic model of both the nucleus and the cytoplasm.

Consider a two-dimensional (2D) cell of thickness *b*_0_, radius *R*_0_ and volume *V*_0_ in its elastic resting state comprising a nucleus of volume *V*_N_ and cytoplasm of volume *V*_C_ such that *V*_0_ = *V*_N_ + *V*_C_. The representative volume element (RVE) of the stress-fibres within the cytoplasm in this resting configuration is assumed to be a cylinder of volume 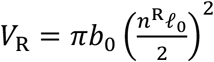, where *ℓ*_0_ is the length of a stress-fibre functional unit in its ground-state, and *n*^R^ is the number of these ground-state functional units within this reference RVE. The total number of functional unit packets within the cell is 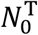, and we introduce 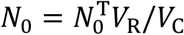 as the average number of functional unit packets available per RVE; *N*_0_ shall serve as a useful normalisation parameter. The state of stress-fibres at location *x_i_* within the cell is described by their angular concentration *η*(*x_i_, φ*), and there are *n*(*x_i_, φ*) functional units in series along the length of each stress-fibre in the RVE. Here, *φ* is the angle of the stress-fibre bundle in the undeformed configuration with respect to the *x*_2_ – direction of the fibronectin stripe. Vigliotti et al. [1] showed that, at steady-state, the number *n*^ss^ of functional units within the stress-fibres is given by

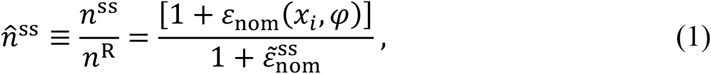

where 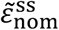 is the strain at steady-state within a functional unit of the stress-fibres, and *ε*_nom_(*x_i_, φ*) is the nominal strain in direction *φ*. The chemical potential of the functional units within the stress-fibres in terms of the Boltzmann constant *k*_B_ is given by

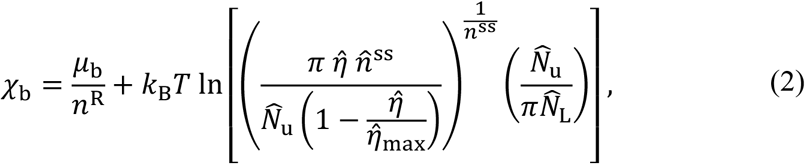

where the normalized concentration of the unbound stress-fibre proteins is 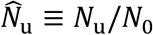 with 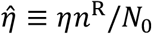, while 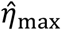 is the maximum normalised value of 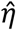 corresponding to full occupancy of all available sites for stress-fibres (in a specific direction) and 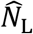 is the number of lattice sites available to unbound proteins. The enthalpy *μ*_b_ of *n*^R^ bound functional units at steady-state is given in terms of the isometric stress-fibre stress *σ*_max_ and the internal energy *μ*_b0_ as

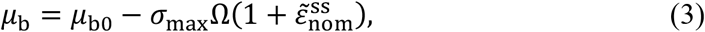

where *Ω* is the volume of *n*^R^ functional units. By contrast, the chemical potential of the unbound proteins is independent of stress and given in terms of the internal energy *μ*_u_ as

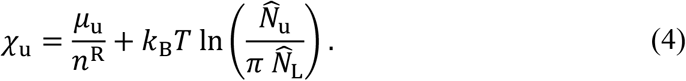

For a fixed configuration of the 2D cell (i.e. a fixed strain distribution *ε*_nom_(*x_i_, φ*)), the contribution to the specific Helmholtz free-energy of the cell *f* from the stress-fibre cytoskeleton follows as

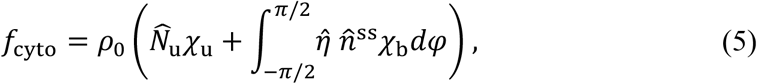

where *ρ*_0_ ≡ *N*_0_/*V*_R_ is the number of protein packets per unit reference volume available to form functional units in the cell. However, we cannot yet evaluate *f*_cyto_ as 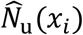 and 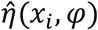 are unknown. These will follow from the chemical equilibrium of the cell as will be discussed subsequently.

The total stress *∑_ij_* within the cell includes contributions from the passive elasticity provided mainly by the intermediate filaments of the cytoskeleton attached to the nuclear and plasma membranes and the microtubules, as well as the active contractile stresses of the stress-fibres. The total Cauchy stress is written in an additive decomposition as

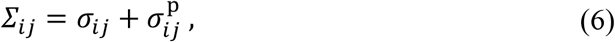

where *σ_ij_* and 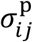 are the active and passive Cauchy stresses, respectively. In the 2D setting with the cell lying in the *x*_1_ – *x*_2_ plane, the active stress is given in terms of the volume fraction 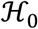 of the stress-fibre proteins as

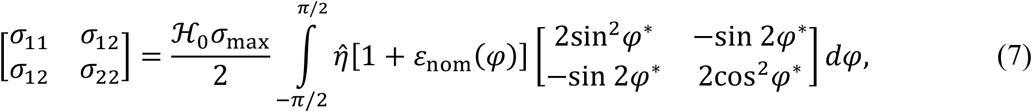

where *φ*^*^ is the angle of the stress-fibre measured with respect to *x*_2_, and is related to its orientation *φ* in the undeformed configuration by the rotation with respect to the undeformed configuration. The passive elasticity in the 2D setting is given by a 2D specialization of the Ogden [2–3] hyperelastic model as derived in [2]. The strain energy density function of this 2D Ogden model is

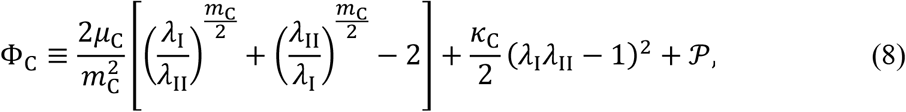

for the cytoplasm and

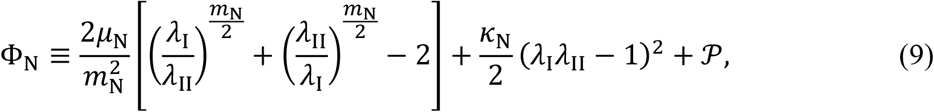

for the nucleus where *λ*_I_ and *λ*_II_ are the principal stretches, *μ*_C_ (*μ*_N_) and *κ*_C_ (*κ*_N_) the shear modulus and in-plane bulk modulus of cytoplasm (nucleus), respectively, while *m*_C_ (*m*_N_) is a material constant governing the non-linearity of the deviatoric elastic response of cytoplasm (nucleus). The term 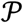 is a penalty term introduced to include an elastic penalty that prevents significant volumetric compression of the cytoplasm/nucleus and given by 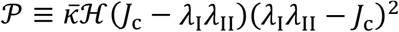, where 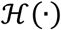 is the Heaviside step function, *J_c_* a non-dimensional constant that sets when this term becomes non-zero and 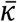 sets the magnitude of the penalty.

The cell is assumed to be incompressible, and thus throughout the cell, we set the principal stretch in the *x*_3_ −direction *λ*_III_ = 1/(*λ*_I_*λ*_II_). The (passive) Cauchy stress then follows as 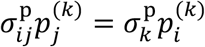 in terms of the principal (passive) Cauchy stresses 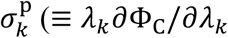 for the cytoplasm and ≡ *λ_k_∂*Φ_N_/*∂λ_k_* for the nucleus) and the unit vectors 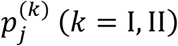 denoting the principal directions. The total specific Helmholtz free-energy of the cell is then *f* = *f*_cyto_ + Φ_C_ in the cytoplasm and *f* = Φ_N_ in the nucleus.

Shishvan et al. [2] have shown that equilibrium of a morphological microstate reduces to two conditions: (i) mechanical equilibrium with *∑_ij,j_* = 0 throughout the system, and (ii) chemical equilibrium such that *χ*_u_(*x_i_*) = *χ*_b_(*x_i_, φ*) = constant, i.e. the chemical potentials of bound and unbound stress-fibre proteins are equal throughout the cell. The condition *χ*_u_ = *χ*_b_ implies that 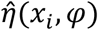 is given in terms of 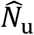, by

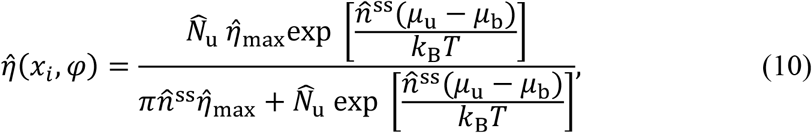

and 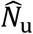 follows from the conservation of stress-fibre proteins throughout the cytoplasm, viz.

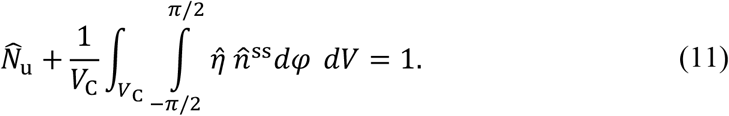

Knowing 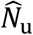 and 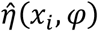, the stress *∑_ij_* can now be evaluated and these stresses within the system (i.e. cell and substrate) need to satisfy mechanical equilibrium, i.e. *∑_ij,j_* = 0. In this case, the mechanical equilibrium condition is readily satisfied as the stress field *∑_ij_* within the cell is equilibrated by a traction field T_*i*_ exerted by the substrate on the cell such that *b∑_ij,j_* = −T_*i*_, where *b*(*x_i_*) is the thickness of the cell in the current configuration. Then *F*_cell_ becomes

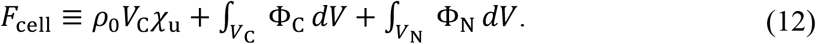

Here, *χ*_u_ is given by Eq. (4) with the equilibrium value of 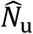 obtained from Eqs. (10–11). For the purposes of further discussion, we label the equilibrium value F_cyto_ ≡ *ρ*_0_V_C_(*χ*_u_ – *χ*_0_) as the cytoskeletal free-energy of the cell where 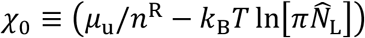 is constant for all microstates and 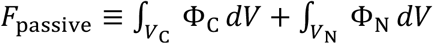 as the passive elastic energy of the cell. The equilibrium value of 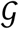 denoted by *G* is then given as *G* = *F*_cell_ + *F*_W_, where 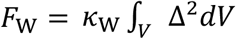 is a numerical energy penalty introduced to prevent the cell exiting the adhesive stripe. The value of Δ is defined as

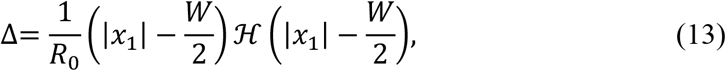

with *W* as the width of the fibronectin stripe. The parameters of the cell free-energy model are listed in Table S 1.

### 2. Numerical methods

#### 2.1 Description of a morphological microstate

The 2D cell in microstate (*j*) is by the *q* positional vectors 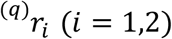. Here we describe the numerical scheme used to denote these vectors. In the 2D context of cells on substrates describing a morphological microstate reduces to specifying the connection of all material points of the cell to locations within the fibronectin stripes, i.e. a displacement field 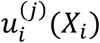 is imposed on the cell with *X_i_* denoting the location of material points on the cell in the undeformed configuration, and these are then displaced to 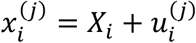 in morphological microstate (*j*), such that all 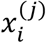 lie within the fibronectin stripe on the micropatterned substrates. These material points located at 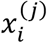 are then connected to material points on the substrate at the same location 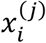, completing the definition of the morphological microstate in the 2D setting.

The cell is modelled as a continuum and thus 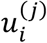 is a continuous field. We define 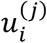 via Non-Uniform Rational B-splines (NURBS) such that the morphological microstate is now defined by *M* pairs of weights 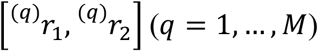. In all the numerical results presented here, we employ *M* = 16 with 4 × 4 weights 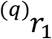 and 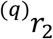 governing the displacements in the *x*_1_ and *x*_2_ directions, respectively. The NURBS employ fourth order base functions for both the *x*_1_ and *x*_2_ directions, and the knots vector included two nodes each with multiplicity four, located at the extrema of the interval. We emphasise here that this choice of representing the morphological microstates imposes restrictions on the morphological microstates that will be considered. Therefore, the choice of the discretization used to represent 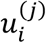 needs to be chosen so as to be able to represent the microstates we wish to sample, e.g. the choice can be based on the minimum width of a filopodium one expects for the given cell type [4]. Given 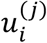, we can calculate *G*^(*j*)^ using the model described above with the cell discretised using constant strain triangles of size *e* ≈ *R*_0_/10.

#### 2.2 Sampling of the homeostatic ensemble

For the results in Fig. S2 where we show distribution of observables under homeostatic conditions, we construct, via MCMC, a Markov chain that serves as a sample of the homeostatic ensemble for cells on substrates. The algorithm closely follows the approach developed by Shishvan et al. [2] and Buskermolen et al. [3] and uses the Metropolis [5] algorithm in an iterative manner using the procedure explained in detail in [2] but now with the following modification for the cells on stripes of width *W*. Typical Markov chains comprised in excess of 20 × 10^6^ samples. The normalized homeostatic temperature 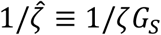, with *ζ* the homeostatic temperature and *G_s_* the Gibbs free energy of the cell in suspended state, obtained from the MCMC simulations is plotted in Fig S2 as a function of the stripe width 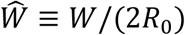 and these results are consistent with those in [3].

#### 2.3 Temporal integration of the HLE

The stationary distributions obtained from the MCMC provided stationary distributions for the observables as well as the homeostatic temperature used in the dimensionless form of the HLE

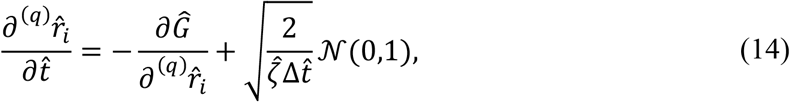

of the HLE. The time integration algorithm is an Euler forwards algorithm, as typically used for stochastic differential equations and is summarised as follows.

i. The cell is placed from suspension on the origin of the substrate at time 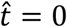.
ii. For the microstate at time 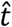 described by the vectors 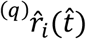 we evaluate the gradient 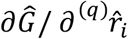 using the function “Derivest” [6]. Derivest is a complex numerical differentiation algorithm that requires 46 separate evaluations of 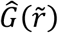; readers are referred to [6] for details of the algorithm.
iii. For all the degrees of freedom we then randomly select the value of the noise from the normal Gaussian distribution, 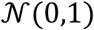 and insert it into (13) to obtain the value of 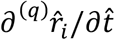.
iv. We then update the current configuration by via the forward Euler scheme.

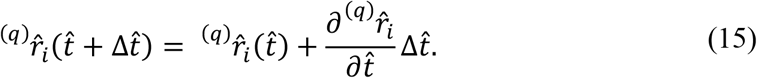
v. All the observables associated to the new cell configuration 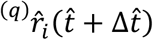 are then stored.
vi. Set 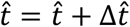, and repeat from step (ii) until 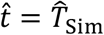. where 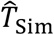 is the maximum simulated time of the experiment.

Time evolutions in both the unpatterned and patterned substrates were computed by selecting 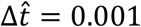.

### 3. Definitions of normalised quantities and observables

Since the cell is modelled as a two-dimensional body changing shape and moving on a micropatterned 2D plane, the typical experimental 2D observables are directly measured from simulations, including morphological observables and protein distributions. The idea is to mimic a live cell imaging time-lapse [7–9] and extract temporal evolution of observables that are representative of measured quantities. This information including the protein distributions can be obtained as a function of time with some examples shown in the Supplementary Videos.

#### 3.1 Morphological observables

The morphological observables describe the cell shape and orientation in the unpatterned or patterned substrate. These are equivalent to analyzing the 2D projection of a single adherent cell on the substrate. The observables of interest, as shown in Fig. S3, are: cell spread area *A*, cell aspect ratio *A_S_*, orientation *ϕ* and cell form factor *FF*. Note that all of these observables are measured as a function of time.

Cell spread area *A* is calculated as the enclosed area within the outer boundary of the cell. The cell aspect ratio *A_S_* and orientation are defined from drawing the best fit ellipse of a given cell configuration. Note that cells take configurations that are not ellipses, but a best fit ellipse provides easy metrics to define cell approximate elongation and orientation and is extensively used in experiments [2–3, 10–12]. The best fit ellipse is calculated as an ellipse that best fits the outline of the current cell configuration. The cell aspect ratio is given by the ratio of the major to minor axis of the best-fit ellipse and the cell orientation is the angle the major axis makes with the stripe direction. The cell form factor FF ≡ *p*^2^/(4*πA*) where *p* is the cell perimiter, is a metric typically used to describe the elongation and capability of the cell to form filopodia on a given substrate. FF depends on the area, but gives completely different information: FF ≫ 1 means that the cell is circular in shape while FF ≫ 1 for a starlike cell, forming many filopodia to sense and explore the environment. The perimeter *p* is calculated as the perimeter of the polygonal shape drawn by the nodes on the periphery of the current cell configuration.

#### 3.2 Protein stainings

The predictions of cell configurations throughout the papers as well as in the Supplementary Videos, show immunofluorescence-like images comprising of the nucleus, actin and vinculin stained in blue, green and magenta, respectively (Fig. S3). Nucleus elements are simply determined from coloring the mesh elements labelled as “nucleus” in blue.

Actin stainings are representations of the stress-fiber distributions explicitly modelled in the framework. These distributions are plotted in order to convey two crucial pieces of information: (i) the concentration of bound stress-fiber units in a given location within the cell configuration (ii) the local orientation of the densest stress-fiber bundle. While the formulation is a continuum formulation the information on stress-fibers is still presented via internal state variables as follows. The concentration of stress-fibers 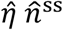 at each location *x_i_* and at an angle *φ*^*^ with respect to the *x*_2_ −axis is known for any cell configuration. We then sketch an actin fibre in the direction 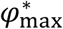 of the maximum value of 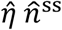 at that location at a spacing that scales inversely with

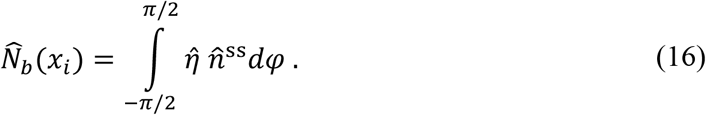

Focal adhesions are not explicitly modelled in the simulations. However, it is well known that mature and larger focal adhesions apply larger tractions on the substrate [13–15]. For these reasons the magnitudes of the traction distribution *T*(*x_i_*) are used as surrogates to visualize the localization of the focal adhesions. Specifically, higher traction magnitudes are represented by a deeper magenta color to visualize suggest adhesion protein density in the given location.

**Table 1:** Model parameters representative of myofibroblasts [3].

| Parameter | Value [units] | Description |
| --- | --- | --- |
| $T$ | 310 [K] | Cell temperature (37°C) |
| $\mu_u = \mu_{u0}$ | 8 kBT | Unbound motor protein reference potential |
| $\mu_{b0}$ | 9 kBT | Bound motor protein reference chemical potential |
| $\sigma_{\max}$ | 240 [kPa] | Maximum stress-fibre tension |
| $\eta_{\max}$ | 0.75 | Maximum allowed angular stress-fibre density |
| $\rho_0$ | $3 \times 10^6$ [packets*$\mu\text{m}^{-3}$] | Density of protein packets that comprise stress-fibres |
| $\mu_{\text{cyto}}$ | 1.67 [kPa] | Cytoplasm shear modulus |
| $m_{\text{cyto}}$ | 5 | Cytoplasm shear exponent |
| $\kappa_{\text{cyto}}$ | 35 [kPa] | Cytoplasm bulk modulus |
| $\mu_{\text{nuc}}$ | 3.3 [kPa] | Nucleus shear modulus |
| $m_{\text{nuc}}$ | 20 | Nucleus shear exponent |
| $\kappa_{\text{nuc}}$ | 35 [kPa] | Nucleus bulk modulus |
| $b_0/R_0$ | 0.2 | Ratio of the resting cell thickness $b_0$ and cell radius $R_0$ in the elastic resting state of the cell |
| $R_N/R_0$ | $0.256\sqrt{\pi}$ | Ratio of the nucleus radius, $R_N$ and $R_0$ |
| $\mathcal{H}_0$ | 0.032 | Volume fraction of stress-fibre proteins |
| $\Omega$ | $10^{-7.1}$ [$\mu\text{m}^3$] | Volume of the reference number of functional stress-fibre units |
| $\bar{\kappa}$ | $10^5$ [kPa] | Numerical penalty modulus |
| $J_c$ | 0.6 | Critical value at which the elastic penalty starts to operate |
| $\kappa_W$ | $5 \times 10^3$ [nN*$\mu\text{m}^{-2}$] | Numerical penalty introduced to maintain cells inside the stripe |

### Supplementary Figures

**Fig. S1:**
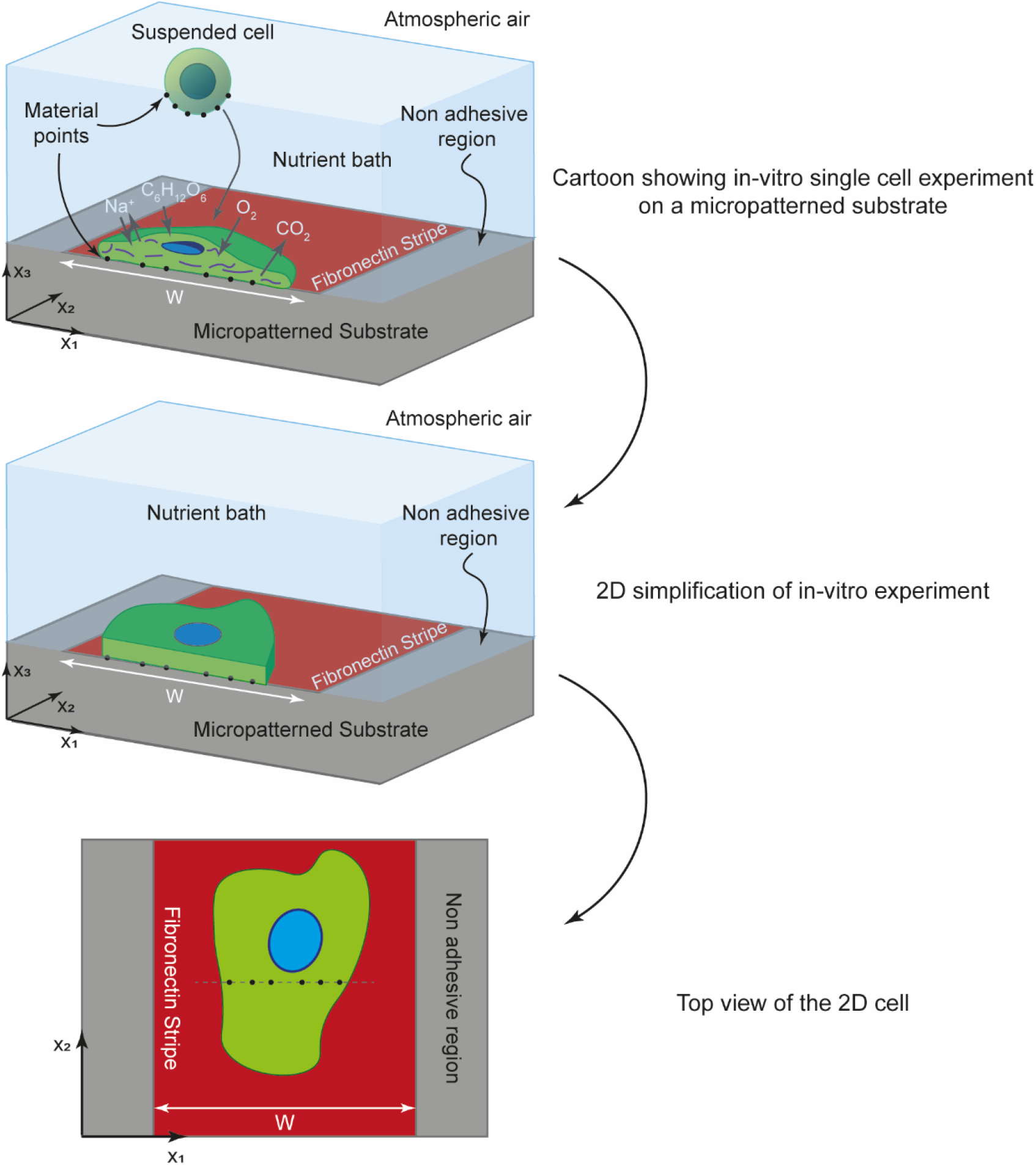
Simplification process of a cell from in-vitro to a 2 dimensional body lying on the *x*_1_ – *x*_2_ plane.

**Fig. S2:**
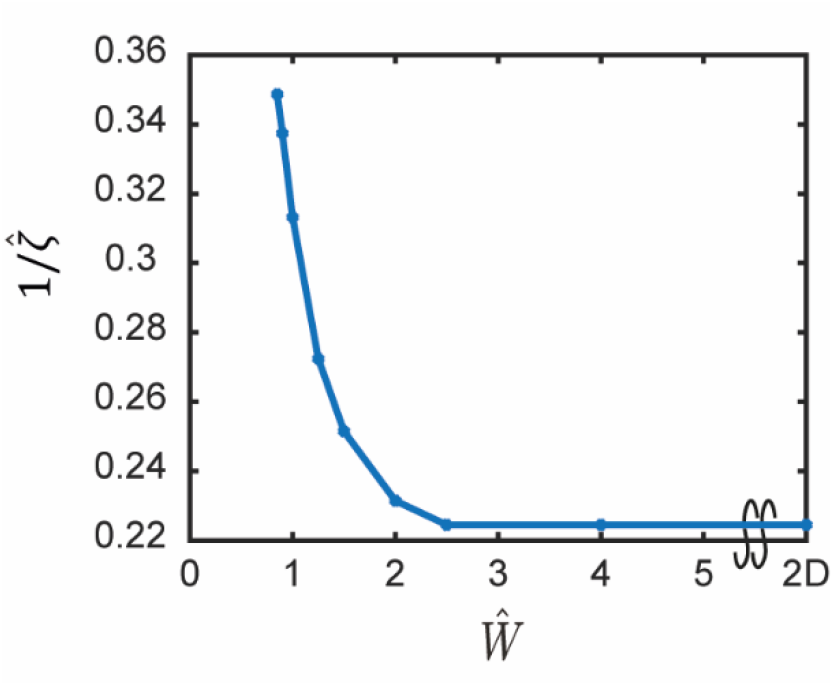
Homeostatic temperature, 1/*ζ*, as a function of fibronectin stripe width, *W*. The homeostatic temperature and linewidth are plotted as the normalized quantities: 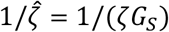 and 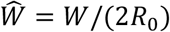.

**Fig. S3:**
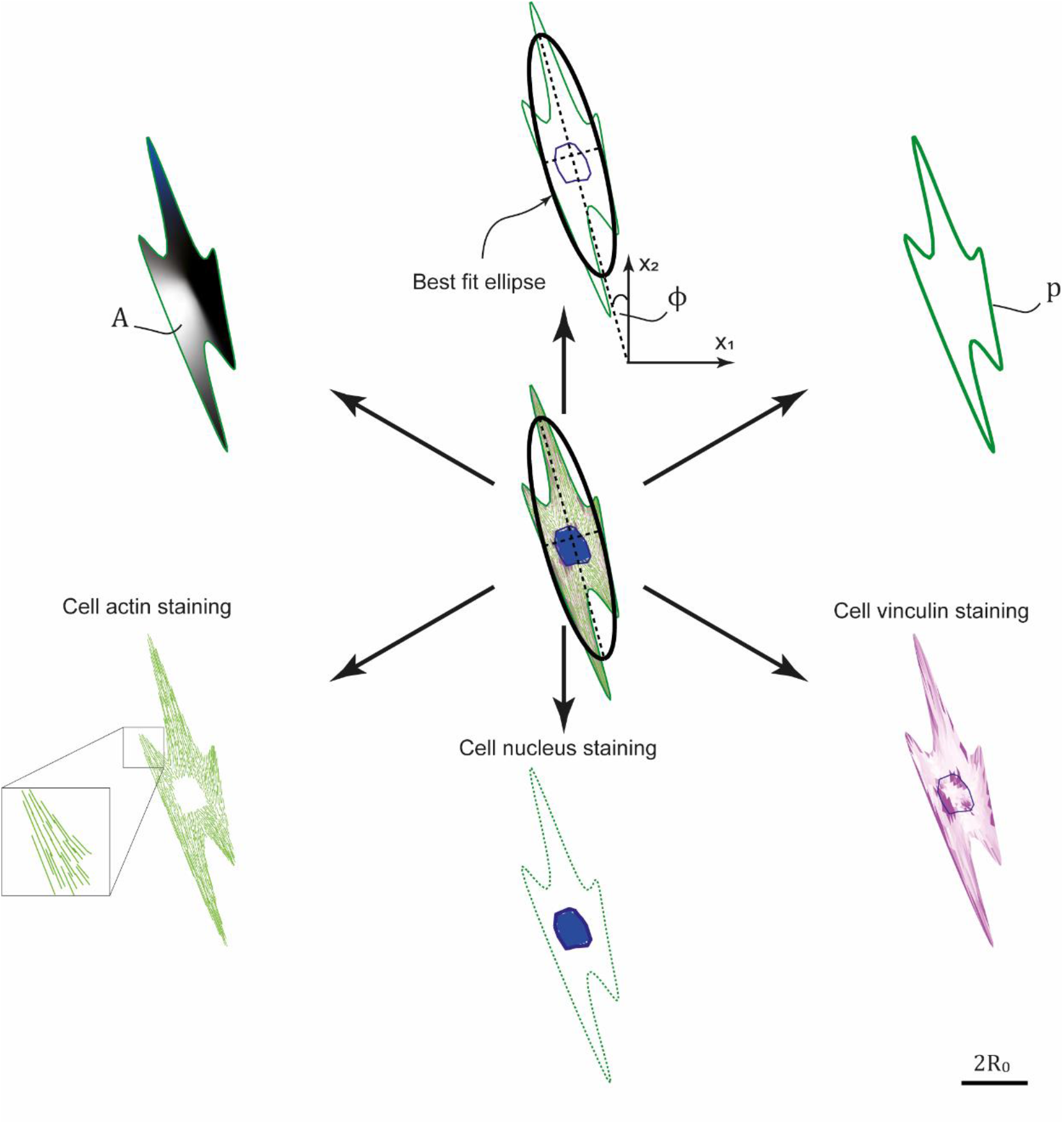
Deriving cell morphological observables and staining distributions from a given cell configuration. Scalebar is 2*R*_0_.

### Supplementary Video Captions

**Supplementary Video 1:** Morphological evolution of a cell. The time-lapse shows the evolution of a cell over a single simulated HLE trajectory after seeding from suspension at normalized time 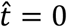. The corresponding evolution of the ensembled averaged morphological observables (normalized area *Â*, aspect ratio *A_s_* and form factor FF) are also included along with an indicator showing the time corresponding to the image of the cell in the movie.

**Supplementary Video 2:** Motility of a cell. The time-lapse shows the coupled motility and morphological evolution of a cell over a single simulation HLE trajectory after seeding from suspension at normalized time 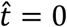. The corresponding temporal evolution of the normalized squared displacement 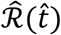 is also included along with an indicator showing the time corresponding to the image of the cell in the movie. The red line shows the path of the centroid of the cell.

**Supplementary Video 3:** Contact guidance of cells. The time-lapses show the coupled motility and morphological evolution of the cells over a single simulation HLE trajectory after seeding from suspension at normalized time 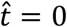 on stripes of two different widths *W* in addition to the 2D substrate where *W* = ∞. The red line shows the path of the centroids of each cell.

## Footnotes

* See Supplementary Section 3.2 for a detailed description of how the model predictions are used immunofluorescence-like images.

## References

1. Aschoff J. Circadian rhythms: influences of internal and external factors on the period measured in constant conditions 1. Zeitschrift für Tierpsychologie. 1979 Jan 12;49(3):225–49.

2. Ohata, K., Nishiyama, H. & Tsukahara, Y. in Biological Clocks: Mechanisms and Applications (ed. Touitou, Y.) 167–170 (Elsevier, Amsterdam, 1999)

3. Johnson DG, Walker CL. Cyclins and cell cycle checkpoints. Annual review of pharmacology and toxicology. 1999;39.

4. Kastan MB, Bartek J. Cell-cycle checkpoints and cancer. Nature. 2004 Nov;432(7015):316–23.

5. Stanewsky R, Kaneko M, Emery P, Beretta B, Wager-Smith K, Kay SA, Rosbash M, Hall JC. The cryb mutation identifies cryptochrome as a circadian photoreceptor in Drosophila. Cell. 1998 Nov 25;95(5):681–92.

6. Kume K, Zylka MJ, Sriram S, Shearman LP, Weaver DR, Jin X, Maywood ES, Hastings MH, Reppert SM. mCRY1 and mCRY2 are essential components of the negative limb of the circadian clock feedback loop. Cell. 1999 Jul 23;98(2):193–205.

7. Kesavan SV, Momey F, Cioni O, David-Watine B, Dubrulle N, Shorte S, Sulpice E, Freida D, Chalmond B, Dinten JM, Gidrol X. High-throughput monitoring of major cell functions by means of lensfree video microscopy. Scientific reports. 2014 Aug 6;4(1):1–1.

8. Nisenholz N, Rajendran K, Dang Q, Chen H, Kemkemer R, Krishnan R, Zemel A. Active mechanics and dynamics of cell spreading on elastic substrates. Soft matter. 2014;10(37):7234–46.

9. Balaban NQ, Schwarz US, Riveline D, Goichberg P, Tzur G, Sabanay I, Mahalu D, Safran S, Bershadsky A, Addadi L, Geiger B. Force and focal adhesion assembly: a close relationship studied using elastic micropatterned substrates. Nature cell biology. 2001 May;3(5):466–72.

10. Bell GI, Dembo MI, Bongrand PI. Cell adhesion. Competition between nonspecific repulsion and specific bonding. Biophysical journal. 1984 Jun 1;45(6):1051–64.

11. Hadden WJ, Young JL, Holle AW, McFetridge ML, Kim DY, Wijesinghe P, Taylor-Weiner H, Wen JH, Lee AR, Bieback K, Vo BN. Stem cell migration and mechanotransduction on linear stiffness gradient hydrogels. Proceedings of the National Academy of Sciences. 2017 May 30;114(22):5647–52.

12. Hadden WJ, Young JL, Holle AW, McFetridge ML, Kim DY, Wijesinghe P, Taylor-Weiner H, Wen JH, Lee AR, Bieback K, Vo BN. Stem cell migration and mechanotransduction on linear stiffness gradient hydrogels. Proceedings of the National Academy of Sciences. 2017 May 30;114(22):5647–52.

13. Engler A, Bacakova L, Newman C, Hategan A, Griffin M, Discher D. Substrate compliance versus ligand density in cell on gel responses. Biophysical journal. 2004 Jan 1;86(1):617–28.

14. Carter SB. Haptotaxis and the mechanism of cell motility. Nature. 1967 Jan;213(5073):256–60.

15. Carter SB. Principles of cell motility: the direction of cell movement and cancer invasion. Nature. 1965 Dec;208(5016):1183–7.

16. Buskermolen AB, Ristori T, Mostert D, van Turnhout MC, Shishvan SS, Loerakker S, Kurniawan NA, Deshpande VS, Bouten CV. Cellular contact guidance emerges from gap avoidance. Cell Reports Physical Science. 2020 May 20;1(5): 100055.

17. Chang SS, Guo WH, Kim Y, Wang YL. Guidance of cell migration by substrate dimension. Biophysical journal. 2013 Jan 22;104(2):313–21.

18. Ray A, Lee O, Win Z, Edwards RM, Alford PW, Kim DH, Provenzano PP. Anisotropic forces from spatially constrained focal adhesions mediate contact guidance directed cell migration. Nature communications. 2017 Apr 12;8(1):1–7.

19. Asano S, Ito S, Takahashi K, Furuya K, Kondo M, Sokabe M, Hasegawa Y. Matrix stiffness regulates migration of human lung fibroblasts. Physiological reports. 2017 May;5(9):e13281.

20. Ghibaudo M, Saez A, Trichet L, Xayaphoummine A, Browaeys J, Silberzan P, Buguin A, Ladoux B. Traction forces and rigidity sensing regulate cell functions. Soft Matter. 2008;4(9):1836–43.

21. Pelham RJ, Wang YL. Cell locomotion and focal adhesions are regulated by substrate flexibility. Proceedings of the National Academy of Sciences. 1997 Dec 9;94(25):13661–5.

22. Guido S, Tranquillo RT. A methodology for the systematic and quantitative study of cell contact guidance in oriented collagen gels. Correlation of fibroblast orientation and gel birefringence. Journal of cell science. 1993 Jun 1;105(2):317–31.

23. Dunn GA, Heath JP. A new hypothesis of contact guidance in tissue cells. Experimental cell research. 1976 Aug 1;101(1):1–4.

24. Weiss P. Experiments on cell and axon orientation in vitro: the role of colloidal exudates in tissue organization. Journal of Experimental Zoology. 1945 Dec;100(3):353–86.

25. Huang CK, Donald A. Revealing the dependence of cell spreading kinetics on its spreading morphology using microcontact printed fibronectin patterns. Journal of The Royal Society Interface. 2015 Jan 6;12(102):20141064.

26. Buskermolen AB, Suresh H, Shishvan SS, Vigliotti A, DeSimone A, Kurniawan NA, Bouten CV, Deshpande VS. Entropic forces drive cellular contact guidance. Biophysical journal. 2019 May 21;116(10):1994–2008.

27. Pathak A, Kumar S. Independent regulation of tumor cell migration by matrix stiffness and confinement. Proceedings of the National Academy of Sciences. 2012 Jun 26;109(26):10334–9.

28. Shishvan SS, Vigliotti A, Deshpande VS. The homeostatic ensemble for cells. Biomechanics and modeling in mechanobiology. 2018 Dec;17(6):1631–62.

29. Peyton SR, Putnam AJ. Extracellular matrix rigidity governs smooth muscle cell motility in a biphasic fashion. Journal of cellular physiology. 2005 Jul;204(1):198–209.

30. Pathak A, Kumar S. From molecular signal activation to locomotion: an integrated, multiscale analysis of cell motility on defined matrices. PloS one. 2011 Mar 31;6(3):e18423.

31. Vigliotti A, McMeeking RM, Deshpande VS. Simulation of the cytoskeletal response of cells on grooved or patterned substrates. Journal of the royal society interface. 2015 Apr 6;12(105):20141320.

32. Shenoy VB, Wang H, Wang X. A chemo-mechanical free-energy-based approach to model durotaxis and extracellular stiffness-dependent contraction and polarization of cells. Interface focus. 2016 Feb 6;6(1):20150067.

33. Bangasser BL, Rosenfeld SS, Odde DJ. Determinants of maximal force transmission in a motor-clutch model of cell traction in a compliant microenvironment. Biophysical journal. 2013 Aug 6;105(3):581–92.

34. Stokes CL, Lauffenburger DA, Williams SK. Migration of individual microvessel endothelial cells: stochastic model and parameter measurement. Journal of cell science. 1991 Jun 1;99(2):419–30.

35. Klank RL, Grunke SA, Bangasser BL, Forster CL, Price MA, Odde TJ, SantaCruz KS, Rosenfeld SS, Canoll P, Turley EA, McCarthy JB. Biphasic dependence of glioma survival and cell migration on CD44 expression level. Cell reports. 2017 Jan 3;18(1):23–31.

36. Gordon., Betts, J. Anatomy and physiology

37. Gibbs JW. Elementary principles in statistical mechanics. Courier Corporation; 2014 Dec 17.

38. Vigliotti A, Ronan W, Baaijens FP, Deshpande VS. A thermodynamically motivated model for stress-fiber reorganization. Biomechanics and modeling in mechanobiology. 2016 Aug;15(4):761–89.

39. Purcell EM. Life at low Reynolds number. American journal of physics. 1977 Jan;45(1):3–11.

40. Alberts B. Molecular biology of the cell.

41. Ichimaru S. Basic principles of plasma physics: a statistical approach. CRC Press; 2018 Mar 8.

42. Plou J, Juste-Lanas Y, Olivares V, Del Amo C, Borau C, García-Aznar JM. From individual to collective 3D cancer dissemination: roles of collagen concentration and TGF-β. Scientific reports. 2018 Aug 24;8(1):1–4.

43. Dunn GA, Brown AF. A unified approach to analysing cell motility. Journal of Cell Science. 1987 Mar;1987(Supplement_8):81–102.

44. Schienbein M, Gruler H. Langevin equation, Fokker-Planck equation and cell migration. Bulletin of Mathematical Biology. 1993 May 1;55(3):585–608.

45. Krummel MF, Bartumeus F, Gérard A. T cell migration, search strategies and mechanisms. Nature Reviews Immunology. 2016 Mar;16(3):193.

46. Lee G, Leech G, Rust MJ, Das M, McGorty RJ, Ross JL, Robertson-Anderson RM. Myosin-driven actin-microtubule networks exhibit self-organized contractile dynamics. Science Advances. 2021 Feb 1;7(6):eabe4334.

47. Recho P, Putelat T, Truskinovsky L. Contraction-driven cell motility. Physical review letters. 2013 Sep 5;111(10):108102.

48. Cardamone L, Laio A, Torre V, Shahapure R, DeSimone A. Cytoskeletal actin networks in motile cells are critically self-organized systems synchronized by mechanical interactions. Proceedings of the National Academy of Sciences. 2011 Aug 23;108(34):13978–83.

## References

1. Vigliotti A, Ronan W, Baaijens FP, Deshpande VS. A thermodynamically motivated model for stress-fiber reorganization. Biomechanics and modeling in mechanobiology. 2016 Aug;15(4):761–89.

2. Shishvan SS, Vigliotti A, Deshpande VS. The homeostatic ensemble for cells. Biomechanics and modeling in mechanobiology. 2018 Dec;17(6):1631–62.

3. Buskermolen AB, Suresh H, Shishvan SS, Vigliotti A, DeSimone A, Kurniawan NA, Bouten CV, Deshpande VS. Entropic forces drive cellular contact guidance. Biophysical journal. 2019 May 21;116(10):1994–2008.

4. Vigliotti A, Shishvan SS, McMeeking RM, Deshpande VS. Response of cells on a dense array of micro-posts. Meccanica. 2020 Jul 17:1–7.

5. Metropolis N, Rosenbluth AW, Rosenbluth MN, Teller AH, Teller E. Equation of state calculations by fast computing machines. The journal of chemical physics. 1953 Jun;21(6):1087–92.

6. D’Errico, J.R., 2007. Derivest.

7. Prasad M, Montell DJ. Cellular and molecular mechanisms of border cell migration analyzed using time-lapse live-cell imaging. Developmental cell. 2007 Jun 5;12(6):997–1005.

8. Pully VV, Lenferink AT, Otto C. Time-lapse Raman imaging of single live lymphocytes. Journal of Raman spectroscopy. 2011 Feb;42(2):167–73.

9. Zattara EE, Turlington KW, Bely AE. Long-term time-lapse live imaging reveals extensive cell migration during annelid regeneration. BMC developmental biology. 2016 Dec;16(1):1–21.

10. Kesavan SV, Momey F, Cioni O, David-Watine B, Dubrulle N, Shorte S, Sulpice E, Freida D, Chalmond B, Dinten JM, Gidrol X. High-throughput monitoring of major cell functions by means of lensfree video microscopy. Scientific reports. 2014 Aug 6;4(1):1–1.

11. Nisenholz N, Rajendran K, Dang Q, Chen H, Kemkemer R, Krishnan R, Zemel A. Active mechanics and dynamics of cell spreading on elastic substrates. Soft matter. 2014;10(37):7234–46.

12. Ghibaudo M, Trichet L, Le Digabel J, Richert A, Hersen P, Ladoux B. Substrate topography induces a crossover from 2D to 3D behavior in fibroblast migration. Biophysical journal. 2009 Jul 8;97(1):357–68.

13. Werner M, Blanquer SB, Haimi SP, Korus G, Dunlop JW, Duda GN, Grijpma DW, Petersen A. Surface curvature differentially regulates stem cell migration and differentiation via altered attachment morphology and nuclear deformation. Advanced science. 2017 Feb;4(2):1600347.

14. Pelham RJ, Wang YL. Cell locomotion and focal adhesions are regulated by substrate flexibility. Proceedings of the National Academy of Sciences. 1997 Dec 9;94(25):13661–5.

15. Frey MT, Tsai IY, Russell TP, Hanks SK, Wang YL. Cellular responses to substrate topography: role of myosin II and focal adhesion kinase. Biophysical journal. 2006 May 15;90(10):3774–82.

